# Degradation of Alzheimer’s amyloid-β by a catalytically inactive insulin-degrading enzyme

**DOI:** 10.1101/2020.04.23.057505

**Authors:** Bikash R. Sahoo, Pritam Kumar Panda, Wenguang Liang, Wei-Jen Tang, Rajeev Ahuja, Ayyalusamy Ramamoorthy

## Abstract

It is known that insulin-degrading-enzyme (IDE) plays a crucial role in the clearance of Alzheimer’s amyloid-β (Aβ). The cysteine-free IDE mutant (cf-E111Q-IDE) is catalytically inactive against insulin, but its effect on Aβ degradation is unknown that would help in the allosteric modulation of the enzyme activity. Herein, the degradation of Aβ(1-40) by cf-E111Q-IDE via a non-chaperone mechanism is demonstrated by NMR and LC-MS, and the aggregation of fragmented peptides is characterized using fluorescence and electron microscopy. cf-E111Q-IDE presented a reduced effect on the aggregation kinetics of Aβ(1-40) when compared with the wild-type IDE. Whereas LC-MS and diffusion ordered NMR spectroscopy revealed the generation of Aβ fragments by both wild-type and cf-E111Q-IDE. The aggregation propensities and the difference in the morphological phenotype of the full-length Aβ(1-40) and its fragments are explained using multi-microseconds molecular dynamics simulations. Notably, our results reveal that zinc binding to Aβ(1-40) inactivates cf-E111Q-IDE’s catalytic function, whereas zinc removal restores its function as evidenced from high-speed AFM, electron microscopy, chromatography, and NMR results. These findings emphasize the catalytic role of cf-E111Q-IDE on Aβ degradation and urge the development of zinc chelators as an alternative therapeutic strategy that switches on/off IDE’s function.

## Introduction

Amyloid-β (Aβ) aggregation is a crucial molecular factor contributing to Alzheimer’s disease (AD) by impairing synaptic function via neuronal cell death.[1,2] Two major Aβ isoforms, Aβ(1-40) and Aβ(1-42), are targeted in controlling AD progression. The anti-amyloid immunotherapy is a promising strategy to control AD progression that minimizes the production of soluble neurotoxic oligomers by rapid fibrillation, redirecting the aggregation pathway to generate amorphous aggregates, and jeopardizing the self-assembly of Aβ monomers by anti-amyloidogenic inhibitors.[3–6] While the modulation of Aβ aggregation by foreign molecules has been a strategic therapy for potential treatment of AD, to the best of our knowledge the effect of anti-amyloidogenic compounds on the proteolysis and clearance of soluble Aβ species has been less explored. Unfortunately, to date, therapeutic advancement has sadly not been effective.

The inability to target Aβ may be attributed to a variety of widely known molecular factors.[7] Although Aβ plaque deposition is moderately correlated to AD, plaque deposition as such has also been identified in healthy brains without dementia.[8,9] Moreover, the involvement of other cellular components (such as membrane constituents, metal ions, enzymes, and apolipoproteins) during Aβ aggregation restricts the strategy for successful therapeutic development.[7,10–14] For example, the interaction of water-soluble Aβ with lipid membrane or metal ions (like Zn^2+^ and Cu^2+^) generates amorphous or low-molecular-weight oligomers that are highly polymorphic and vary in neurotoxicity.[15–19] The metal-bound Aβ species have been established as a stable, toxic, and pathologically important target for AD.[20] Metal ion binding to amyloid peptides can affect their enzymatic degradation.[21–24] Reduction in Aβ accumulation is evidenced by enzymes such as insulin-degrading enzyme (IDE), neprilysin (NEP), endothelin-converting enzyme, and matrix metalloproteinase-9. IDE and NEP are reported to clear soluble Aβ, whereas matrix metalloproteinase-9 enzyme was found to degrade both soluble Aβ species and Aβ fibers.[25] NEP is a membrane-bound protein and its expression in neurons is regulated by γ-secretase that is involved in the processing of amyloid precursor protein (APP).[26] On the other hand, IDE, a conserved Zn^2+^ metallopeptidase, is majorly present in the cytosol with a small fraction in the extracellular space that interacts selectively with Aβ monomers which coexist with metal ions.[27] IDE’s activity is mediated by the dynamic equilibrium between soluble Aβ monomers and aggregates.[28] While degradation/clearance of Alzheimer’s Aβ is essential, it is important to develop strategies to avoid the degradation of functional proteins (for example, insulin). For this purpose, studies have focused on the development of small molecules that are capable of selectively regulating the enzymatic activity of IDE against pathological proteins (for example, Aβ) without affecting functional proteins like insulin. [29–31] For example, simultaneous inhibition of insulin proteolysis and ∼2.5-fold enhancement of Aβ degradation by dynorphin B-9 has been reported.[32] The mechanism and action of small molecules on IDE’s activity have been probed by mutational studies. In fact, allosteric mutation has been proposed to be an important strategy for the selective activation of IDE activity against Aβ.[33] Thus, establishing a correlation between IDE mutation and its catalytic function in particular against Aβ degradation is important, which could guide the development of allosteric modulation of IDE’s activity.[29,33] The IDE mutants, E111Q and cysteine-free E111Q (hereafter cf-E111Q-IDE), have been reported to be inactive against insulin.[34] While E111Q has been shown to inhibit Aβ aggregation based on fluorescence experiments [35], the catalytic activity of these mutants against Aβ has not been well studied. In this study, we report the first comprehensive evaluation on the ability of cf-E111Q-IDE to degrade Aβ *in vitro* by using a combination of different biophysical techniques. We further show that zinc binding to Aβ jeopardizes the catalytic activity of cf-E111Q-IDE, whereas zinc removal from the Zn-Aβ complex resumes the catalytic activity of cf-E111Q-IDE.

## Results

### cf-E111Q-IDE mutant delays Aβ(1-40) fibrillation

The catalytic rates of the wild-type IDE (WT-IDE) and the cysteine-free mutant (cf-E111Q-IDE) co-incubated in the presence and absence of EDTA were tested on the rate of hydrolysis of a bradykinin-mimetic fluorogenic peptide, substrate V.[39] Results showed that WT-IDE was active on substrate V hydrolysis while cf-E111Q-IDE mutant or IDE variants co-incubated with EDTA were not (Figure S1). We next tested the activities of WT-IDE and cf-E111Q-IDE in the absence of EDTA (zinc-bound Aβ) and presence of EDTA (zinc-free Aβ) on Aβ(1-40) using thioflavin T (ThT) fluorescence assay (Figure S2). At sub-stoichiometric IDE concentrations (enzyme:Aβ=1:1000 and 1:250 molar ratios), Aβ aggregation showed WT-no ThT fluorescence in the presence of WT-IDE whereas a small delay (t_1/2_≈9 hours) was observed in the presence of cf-E111Q-IDE as compared to that observed for the enzyme-free Aβ sample (Figure S2). On the other hand, WT-IDE mixed with EDTA showed Aβ aggregation with a substantial delay (t_1/2_≈45 hours) whereas cf-E111Q-IDE-EDTA showed a similar effect as observed in the absence of EDTA (Figure S2). Remarkably, at a higher concentration (1:10 enzyme:Aβ molar ratio), all IDE variants showed no Aβ(1-40) aggregation up to day 12 (Figure S3) [28,35]; fluorescence intensities were collected continuously (with 5 minutes interval) up to 60 hours, and then change in intensity was monitored for every 24 hours until day-12 to measure Aβ aggregation. Zinc-free IDE variants were obtained by pre-incubating the enzyme (that contain bound zinc) with EDTA followed by filtration. Zinc-free IDE showed a substantial catalytic activity as indicated by no Aβ aggregation at 1:10 enzyme:Aβ molar ratio (Figure S3). These results indicate that IDE variants are catalytically active against Aβ(1-40) fibrillation despite selective mutation or zinc chelation above a certain threshold concentration of IDE. It should be mentioned here that the threshold concentration tested in this study (0.5 µM) is against ten molar excess of Aβ(1-40); however, it is evidenced that in the brain, the amount of IDE is higher than Aβ. [40] Our observations of ThT fluorescence quenching by the IDE mutant as shown in Figures S2 and S3 can also be due to a possible chaperone activity of the enzyme as hypothesized in previous studies.[35,41]

### Proteolysis of Aβ(1-40) by cf-E111Q-IDE

To test the above-mentioned hypothesis, we assumed that all variants of IDE generate fragments of Aβ(1-40) under a catalytic environment as reported earlier for WT-IDE.[28] In contrast, under a chaperone environment, IDE is expected to exhibit no change on Aβ(1-40) molecular size or no increase in Aβ size due to aggregation. We performed HPLC-mass spectrometry (LC-MS) analysis to characterize the Aβ(1-40) species incubated with WT-IDE, cf-E111Q-IDE, and cf-E111Q-IDE-EDTA at 1:10 IDE:Aβ molar ratio (Figures S4-S6). Strikingly, LC-MS identified Aβ(1-40) fragments in all these three IDE samples (in washed flow-through, see methods) with molecular weights varying from >200 to <800 Da (Figures 1A-C and S4-S6). Several IDE cleavage sites in Aβ have been reported earlier [28] (Figure 1D) that includes N-terminal (V12-H13, H13-H14, H14-Q15, V18-F19, F19-F20, F20-A21) and C-terminal (K28-G29) sites that produce fragments of size >1 kDa. Several other C-terminal cleavage sites (G33-L34, L34-M35, and M35-V36) have also been identified that generate small fragments of size <1 kDa. Since Aβ(1-40) fragments have a tendency to aggregate, [28] the large-size fragments could have been removed during sample filtration using a 10-kDa filter prior to LC-MS measurement. The observations from LC-MS experiments confirm that WT-IDE, cf-E111Q-IDE, and zinc-free cf-E111Q-IDE are catalytically active against Aβ(1-40), which is in agreement with the ThT fluorescence results. The TEM analysis confirmed the presence of large size globular species in all IDE-Aβ mixed samples with size >20 nm (Figure 1E-H), whereas Aβ(1-40) showed a fibrillary morphology after ∼24 hours of incubation (Figure 1E). The presence of morphologically similar non-fibrillary spherical Aβ species in both wild-type and mutant IDE mixture indicates a similar proteolytic activity in all the three samples. We note that the observed catalytic activity of IDE is specific to Aβ(1-40), but not to substrate-V (Figures 2A and S1). Specifically, the cf-E111Q-IDE mutant and zinc-free IDE were found to be catalytically inactive to hydrolyze the fluorogenic substrate-V, but they were active to degrade Aβ(1-40) as revealed by LC-MS and TEM results.

**Figure 1.**
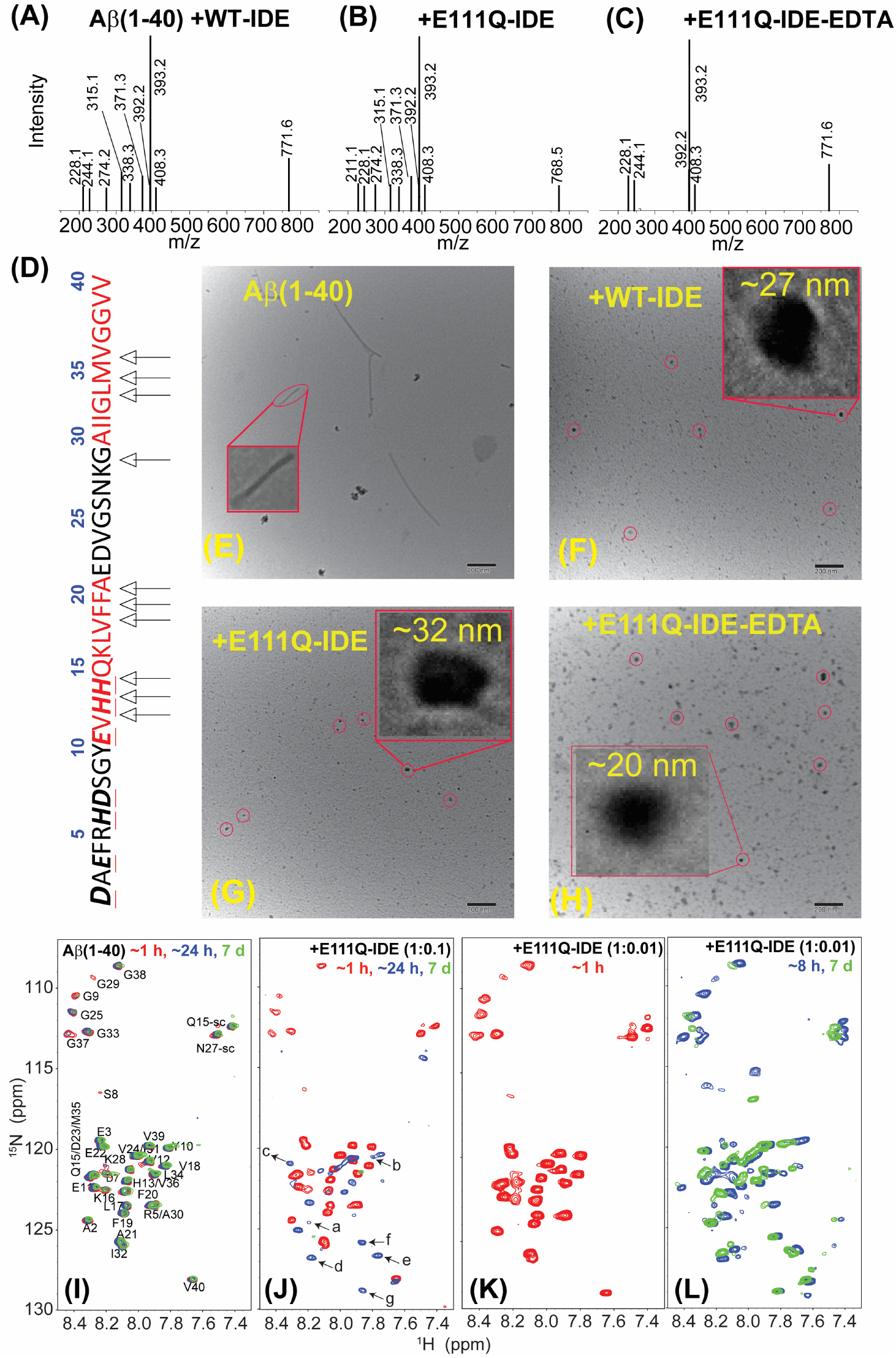
Characterization of Aβ(1-40) species in the presence of IDE. (A-C) HPLC-MS spectra revealing the formation of Aβ fragments by wild-type, E111Q mutant, and zinc-free E111Q-IDE variants as indicated. 50 µM of Aβ(1-40) monomers were incubated with 5 µM of IDE variants for ∼12 hours at room temperature prior to HPLC-MS sample preparation. The original HPLC-MS spectra are shown in supporting Figures S4-S6. (D) Schematic showing Aβ(1-40) amino acids and IDE cleavage sites as indicated by arrows, amino acid residues in the β-strand regions in amyloid fibril are shown in red. The possible zinc-binding residues are underlined. (E-H) Negatively stained TEM images of 5 µM Aβ(1-40) in the absence or presence of 0.5 µM WT-IDE, E111Q-IDE, and E111Q-IDE+5µM EDTA. The samples are incubated ∼24 hours at room temperature prior to imaging. The scale bar is 200 nm. (I-L) 2D SOFAST-HMQC spectra of 25 µM Aβ(1-40) dissolved in 10 mM sodium phosphate (NaPi) buffer, pH 7.4, 10% D_2_O mixed with or without 2.5 or 0.25 µM E111Q-IDE as indicated (non-overlapped NMR spectra are shown in extended figures in the supporting information (Figures S13 and S14)). The new unassigned peaks (denoted as a-g) appeared in (J) ar6e indicated by arrows. NMR spectra were recorded on a 600 MHz spectrometer at 25 °C at different time intervals as indicated. E111Q-IDE (in B, C, G, H, and J-L) indicates the cysteine-free E111Q-IDE mutant.

**Figure 2.**
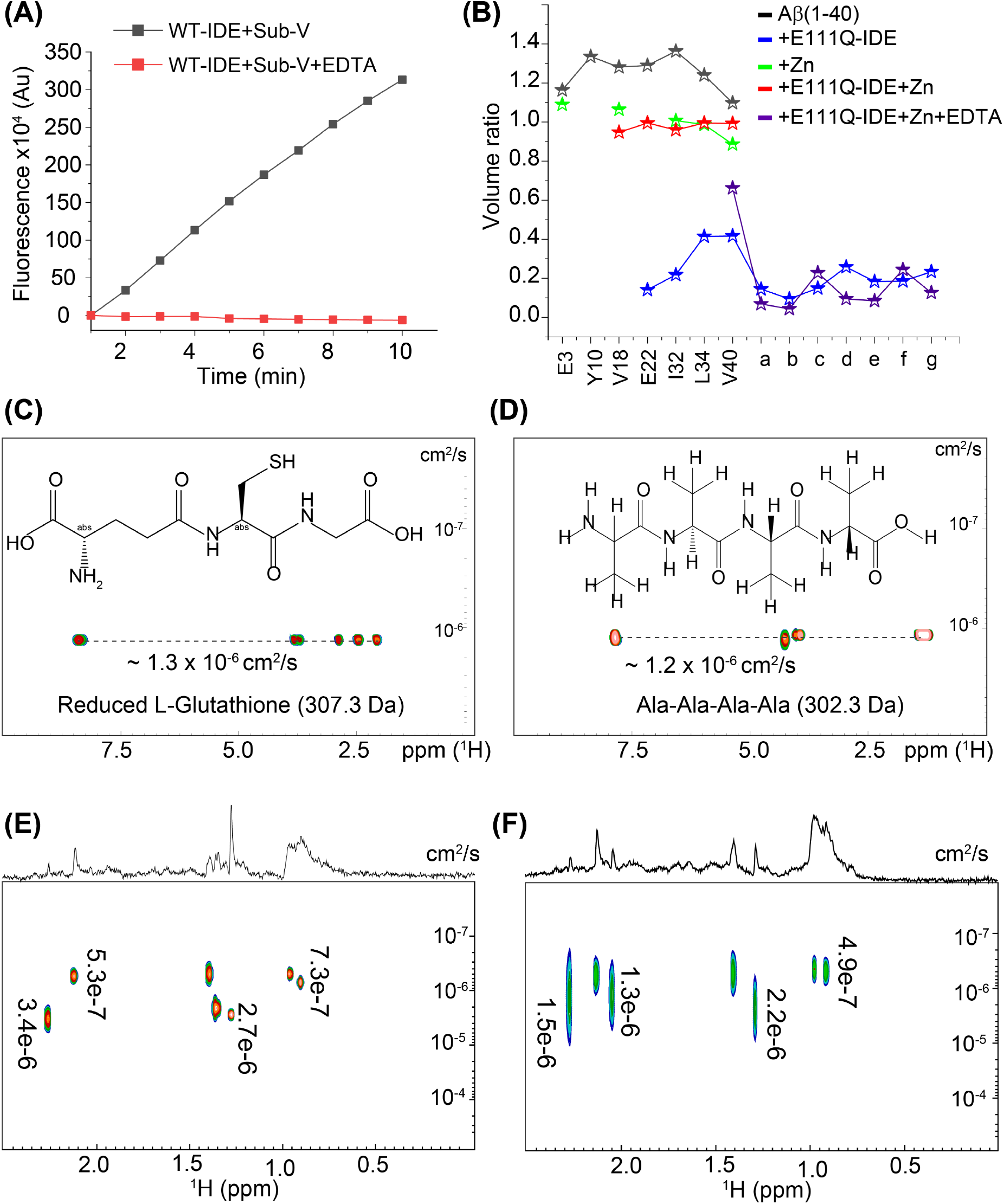
Monitoring the catalytic activity of 0.5µM wild-type IDE incubated with (red) or without (black) equimolar EDTA overnight on Substrate-V (5 µM) as a function of time. (B) Mapping the depletion in Aβ(1-40) residue peak volume for a selected set of non-overlapping ^15^N/^1^H cross-peaks and unassigned newly appeared peaks (denoted as a-g in Figure 1J and 6I) in the absence or presence of zinc, EDTA, and E111Q-IDE as indicated in colors. The plotted peak volumes (volume at ∼24 hours/volume at ∼1 hour) were calculated using the NMR spectra shown in Figures 1 (I, J), 5(E, F), and 6(I). Characterization of wild-type and E111Q-IDE cleaved Aβ(1-40) fragments by DOSY NMR. DOSY NMR spectra of reduced 0.5 mg/mL L-Glutathione (C) and Ala-Ala-Ala-Ala (D). The chemical structure of both peptides and their diffusion constants (cm^2^/s) are indicated inside the graph. 2D DOSY spectrum of 25µM Aβ(1-40) incubated with 2.5 µM of wild-type (E) or E111Q-IDE (F). The corresponding 1D spectrum is shown on the top of the 2D spectrum. Diffusion coefficients are calculated using Bruker Dynamics Center and indicated in each graph. All DOSY NMR samples were prepared in 10 mM NaPi, pH 7.4 containing 10% D_2_O, and spectra were recorded at 25 °C9using a Bruker 500 MHz NMR spectrometer. The cysteine-free IDE mutant is denoted as E111Q-IDE in (B).

### Real-time monitoring of Aβ(1-40) degradation by cf-E111Q-IDE

The ThT fluorescence, TEM, and LC-MS results correlate with each other; however, these results do not provide substantial information about the aggregation of degraded Aβ(1-40) peptide fragments in real-time. To obtain this information, we performed NMR measurements to monitor the enzymatic degradation activity of cf-E111Q-IDE. NMR experiments can be used to reveal the fragmentation of Aβ(1-40) by IDE as the variation in chemical environment and population can alter the frequency, intensity and width of spectral lines. Moreover, the solution NMR method employed here is sensitive to monomers and small-size oligomers, while large oligomers and fibers are not detected due to their large size and slow tumbling rate. As shown in Figure 1I, Aβ(1-40) incubated for ∼1 hour at room temperature presented a well dispersed ^15^N/^1^H correlation spectrum resembling an unstructured conformation as reported previously.[4] A decrease in signal intensity was observed at ∼24 hours and 7-day incubation (Figure 1I). Remarkably, Aβ(1-40) incubated with cf-E111Q-IDE at a stoichiometry of 1:10 enzyme:substrate showed no major changes in the spectrum at ∼1 hour, but several new peaks appeared at ∼24 hours of incubation (Figure 1J, blue). The peak volume analysis of selected non-overlapping residues, as shown in Figure 2B, presented a substantial decrease in signal intensity for the original peaks and the new peaks showed a volume ratio of ∼10-20% in the spectrum of cf-E111Q-IDE-Aβ(1-40) mixture after ∼24 hours of incubation. This observation indicates the formation of Aβ peptide fragments, which has previously been observed for Aβ in the presence of WT-IDE.[28] Strikingly, most of the peaks in the spectrum disappeared on day-7 (Figure 1J, green) suggesting the formation of large-size aggregates by Aβ(1-40) fragments that are beyond the detection limit of solution NMR spectroscopy. This observation correlates well with the TEM observations that showed particles of size ∼32 nm (Figure 1G).

Next, we used NMR experiments to monitor the cleavage of Aβ(1-40) by cf-E111Q-IDE at a sub-stoichiometric concentration (enzyme:substrate=1:100 molar ratio) to ensure the concentration-dependent proteolytic activity of the enzyme as evidenced from the ThT fluorescence (Figure S2). The 2D ^15^N/^1^H chemical shift correlation spectrum showed the generation of Aβ(1-40) fragments with the appearance of several new peaks (Figure 1K, L), but comparatively a slow degradation was observed until day-7. Importantly, on day-7, unlike our observation at a higher concentration (1:10 enzyme:Aβ) of cf-E111Q-IDE (Figure 1J, green), we did not observe a significant signal loss at low (1:100 enzyme:Aβ) concentration (Figure 1L, green). This could be explained as the kinetics of Aβ(1-40) self-assembly to form NMR visible lower-order aggregates is faster than the fragmentation of monomers by cf-E111Q-IDE at low cf-E111Q-IDE concentration. Considering the proteolytic cavity of IDE (volume ∼1.3×10^4^ Å^3^) as evidenced from the X-ray structure,[36] the lower-ordered NMR visible Aβ aggregates that do not fit the active site cannot be the substrates of cf-E111Q-IDE. Therefore, the NMR data (Figure 1L, green) suggests that the Aβ(1-40) oligomers appearing on day-7 are not the substrates of cf-E111Q-IDE and are likely to be structurally different from that of monomers.

### NMR detection of short Aβ(1-40) fragments cleaved by cf-E111Q-IDE

Next, we performed DOSY based NMR experiments to estimate the size of the Aβ(1-40)-fragments generated by cf-E111Q-IDE. For reference, we first calculated the diffusion constants of two short peptides, namely Ala-Ala-Ala-Ala (302.3 Da) and reduced L-Glutathione (307.3 Da), under identical experimental conditions. The diffusion constants of Ala-Ala-Ala-Ala and reduced L-glutathione were calculated as ∼1.2×10^−6^ and ∼1.3×10^−6^ cm^2^/s, respectively, in 10 mM NaPi, pH 7.4 containing 10% D_2_O (Figure 2C and D). WT-The 1:10 WT-IDE:Aβ molar ratio sample incubated for ∼3 hours presented a variable size NMR detectable species with diffusion constant ranging from 3.4×10^−6^ and 7.3×10^−7^ cm^2^/s(Figures 2E and S7A). Similar to the WT-IDE:Aβ sample, several Aβ(1-40) fragments were also identified in the presence of cf-E111Q-IDE at ∼3 hours with the diffusion constant values varying between 1.5×10^−6^ and 4.9×10^−7^ cm^2^/s (Figures 2E, F and S7). The diffusion constants measured from the two representative peaks at 0.90 and 1.27 ppm of the Aβ-cf-E111Q-IDE sample are 2.2×10^−6^ and 6.5×10^−7^ cm^2^/s, respectively (Figure S7B). The diffusion value obtained from peak 0.90 ppm is nearly equal to that for the short peptides Ala-Ala-Ala-Ala and L-Glutathione. This finding indicates that the degradation of Aβ by cf-E111Q-IDE is similar to that of the WT-IDE. The detection of these peptide fragments at ∼24 hours by NMR indicates their water solubility (Figure S7), and correlates to our ^15^N/^1^H SOFAST HMQC observation at ∼24 hours (Figure 1J). Thus, the DOSY results confirmed that both WT-IDE and cf-E111Q-IDE are able to proteolytically cleave Aβ(1-40) to generate variable size fragments. Taken together, the NMR findings, in combination with other biophysical results, highlighted the proteolytic activity of cf-E111Q-IDE to modulate Aβ(1-40) aggregation that was previously hypothesized to be mediated through a non-proteolytic pathway.[35]

### Comparative structural models for full-length Aβ(1-40) and IDE cleaved Aβ aggregates obtained from atomistic simulation

The adeptness of aggregation of Aβ(1-40) fragments generated by IDE and its correlation with morphologically distinct toxic phenotypes were next investigated using explicit solvent all-atom MD simulations. We have compared the self-assembly efficacy of a mostly unstructured (Figure S8) full-length Aβ(1-40) with 20 different IDE-cleaved fragments (Figure 3A) that were previously identified using mass spectrometry. These Aβ fragments (except 1-12 and 36-40) can be further degraded by IDE into smaller fragments as shown in Figure 1D, but were not considered for the simulation studies. The residue-specific solvent accessible surface area (SASA) values are shown in Figure 3A. The SASA values were calculated for all the non-solvent atoms that include the 9 copies of peptide monomers which exhibited intermolecular association during MD simulation. The simulation results identified two trends indicated by blue (high SASA values) and orange (low SASA values) in Figure 3A during the 500 ns MD simulation. Among the 20 peptide fragments, the N-terminal (1-12, 1-13, 1-14, 1-18 and 1-19) and C-terminal (34-40, 35-40 and 36-40) fragments showed relatively high SASA values as compared to the full-length Aβ(1-40) (Figure 3A). While it is not surprising to see the high SASA values for the N-terminal hydrophilic fragments, the high SASA values for the hydrophobic C-terminal fragments may be attributed to their short lengths which is further supported by their no aggregation propensity during the 500 ns MD simulation.

**Figure 3.**
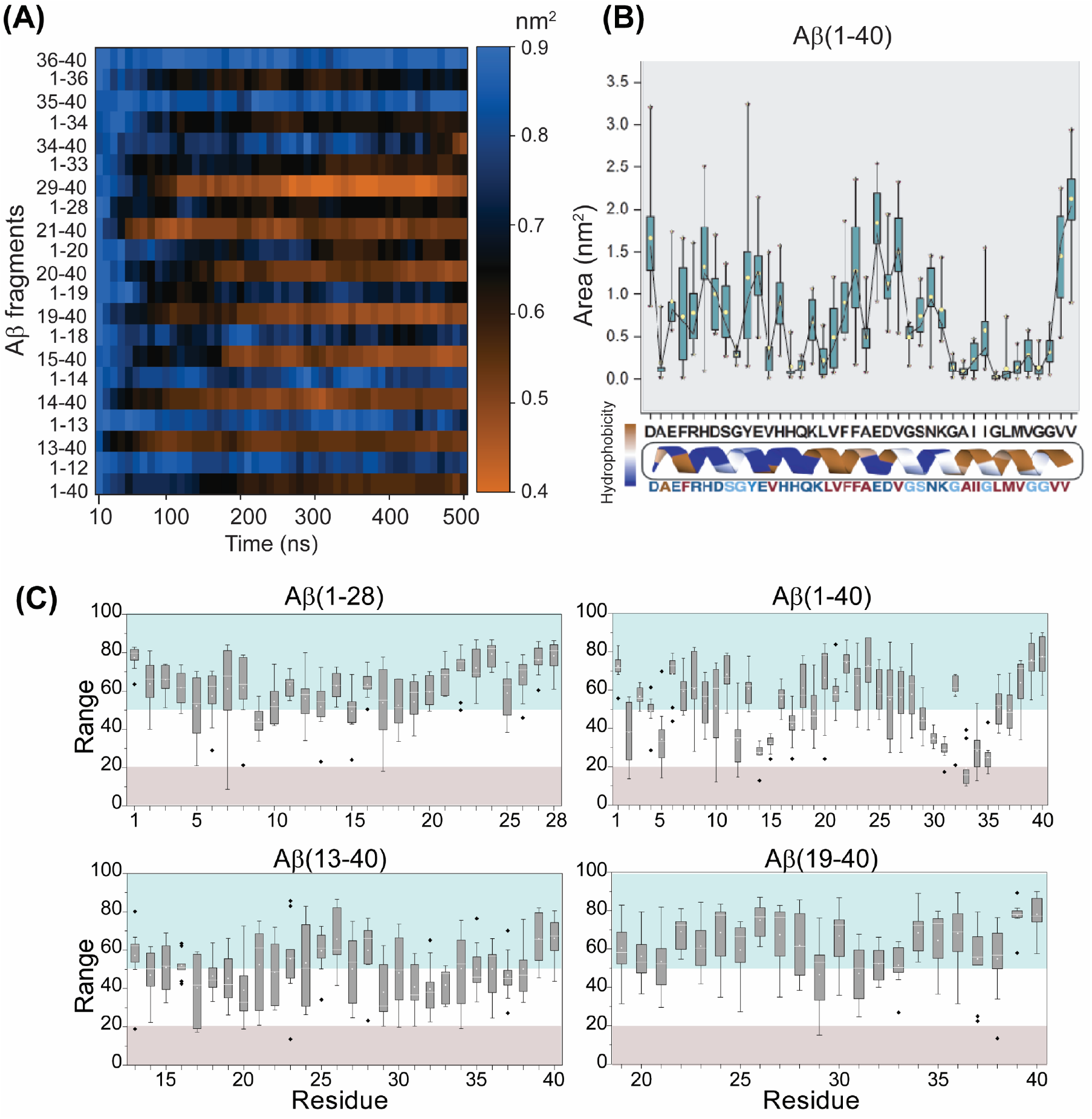
Atomistic simulation of the full-length Aβ(1-40) obtained from 2µs MD simulation (see Figure S8) and for the 20 different IDE cleaved fragments. (A) The SASA values for Aβ fragment atoms accessible to water within a radius of 1.4 Å during the course of 500ns simulation. Blue depicts higher SASA and orange depicts lower SASA as indicated on the scale bar. (B) Average area of full length Aβ(1-40) as a function of residues calculated from the 500ns MD simulation. The dotted line show residues with an average area below 1 nm^2^. (C) Box plot showing the residue-specific solvent exposure of full-length different sized Aβ fragments as indicated. The range (0-100) represents the solvent exposed (>50) and buried residues (<20) in the Aβ fragments. The grey illustrates buried residues, and blue shows solvent exposed residues. The range shown in the plot are the average values calculated from 5 structures derived at equal time-intervals from the last 200 ns MD simulation. The standard error and median (–) are plotted for individual residue.

Self-assembling of Aβ fragments was also measured by calculating the residue-specific average area of the nine molecules initially placed ∼0.5 nm away from each another. A large area indicates the non-aggregated peptide species in an aqueous medium whereas a small area is an indication of an aggregated species. As shown in Figure 3B, a majority of Aβ(1-40) residues presented an average residue-specific area of ≤1.0 nm^2^ indicating self-assembling during the 500 ns MD simulation. Long Aβ fragments such as Aβ(13-40), Aβ(21-40) and Aβ(1-33) also presented a small (≤1.0 nm^2^ per residue) area suggesting their aggregation, whereas short fragments such as Aβ(1-12) and Aβ(35-40) were found to have a large area (≥1.5 nm^2^) as shown in Figure S9.

The neurotoxicity of Aβ oligomers is likely because of an exposed hydrophobic surface spanning residues 17-28 which is facilitated by the concurrent shielding of the charged N-terminus residues 1-12 as previously proposed.[42] To validate this molecular feature and to explain the distinct morphological toxic phenotypes of the full-length versus the IDE-fragmented peptides, we compared the SASA values and the simulated structures of the full-length Aβ(1-40) and IDE-cleaved Aβ-fragments (Figures 3C). Our results obtained from the full-length Aβ(1-40) MD simulation showed that the central hydrophobic residues 21-28 are solvent exposed whereas several N- and C-terminal residues are found to be buried (Figure 3C, grey). Interestingly, the hydrophobic residues 21-28 are solvent exposed in 1-28 and 19-40 fragments, whereas they are buried in the 13-40 fragment (Figure 3C). These observations indicate that the hydrophilic 1-12 residues are important in protecting the solvent exposure of the central hydrophobic core regions that have been proposed to enhance the toxic feature of Aβ oligomers. Based on these results, we speculate that the deletion of N-terminal 1-12 residues by IDE could explain the non-toxic nature of the IDE-fragmented Aβ aggregates.[28,35]

Structural mapping of the full-length Aβ(1-40) peptide and the long fragments, including Aβ(13-40) and Aβ(21-40), identified a mixture of monomers, dimers, trimers, and tetramers from the 500 ns MD simulation (Figure 4(A, C and D)). In contrast, both N and C-terminal fragments, including Aβ(1-12) and Aβ(35-40), showed a distribution of monomers or dimers majorly during the 500 ns MD simulations (Figure 4(B and E)). These findings explain the observation of detectable small species in DOSY NMR and their non-aggregating properties (Figure 2F). Overall, the simulation results pictured distinct self-assembling features of the IDE cleaved Aβ fragments studied here; however, in a natural environment, hetero-assembly of Aβ fragments can also be possible, which was not probed in the present study.

**Figure 4.**
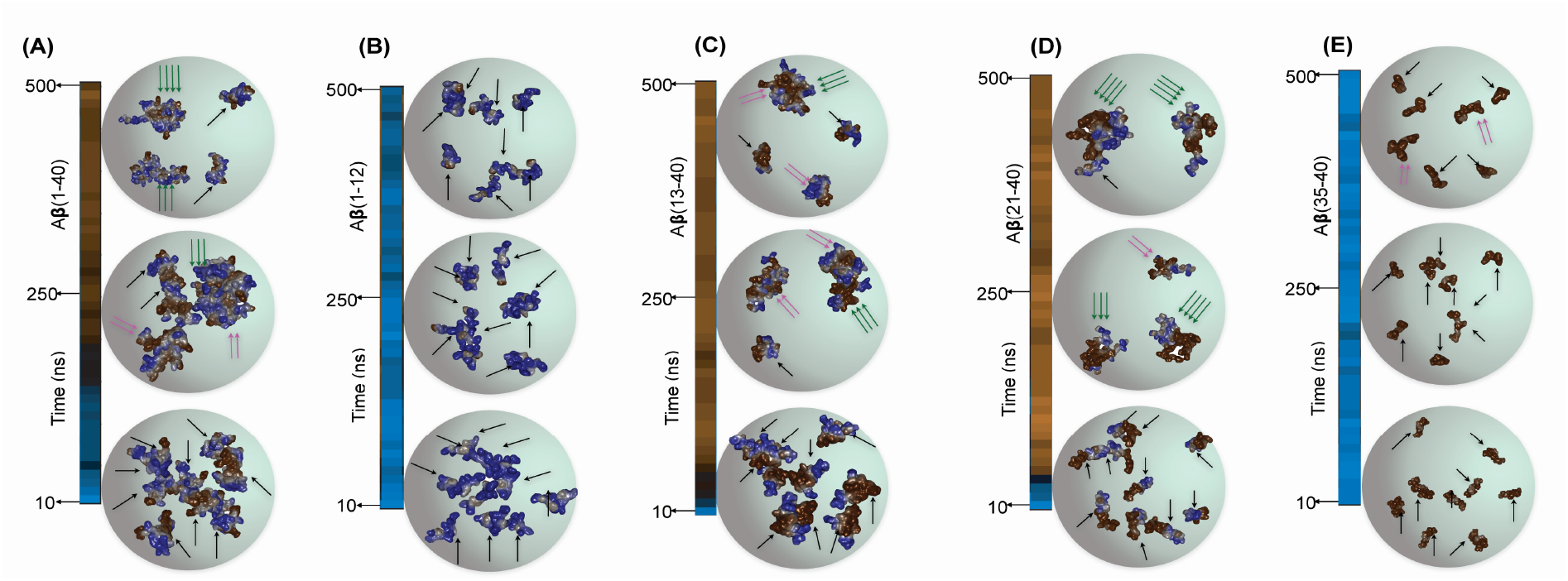
Atomic resolution structures of (A) Aβ(1-40) and its fragments (B-E) were derived from the 500 ns MD simulation and illustrating the structures derived at three different time points (10, 250 and 500 ns). The arrows represent the peptide monomer units in the self-assembled aggregates: a single black arrow for monomers, two pink arrows for dimers and three green arrows for trimers or tetramers.

### Effect of zinc on Aβ(1-40) proteolysis by cf-E111Q-IDE

Following NMR observation, that showed size-dependent proteolytic activity of cf-E111Q-IDE, we next investigated the effect of cf-E111Q-IDE on the zinc-Aβ(1-40) complex that has been shown to be toxic and prevents Aβ-degradation by NEP, IDE, and matrix metalloprotease. [24,43,44] As shown in Figures 5A and S10, the ThT fluorescence shows retardation and acceleration of Aβ(1-40) aggregation by zinc and EDTA, respectively. It is remarkable that both zinc-bound and zinc-free cf-E111Q-IDE are capable of suppressing Aβ aggregation as evidenced from ThT fluorescence (Figure 5A) and NMR (Figure S11A) results. These observations indicate that zinc-chelated-Aβ(1-40) could be a substrate of cf-E111Q-IDE. To investigate the effect of zinc binding to Aβ(1-40) and its removal from the zinc-Aβ(1-40) complex, ^1^H NMR experiments were carried out. A substantial reduction in signal intensity in 6.5 to 8.5 ppm region of the Aβ(1-40) was observed when mixed with equimolar zinc, indicating the formation of zinc-peptide aggregates (Figure 5B). Remarkably, this zinc binding induced signal intensity reduction was mostly recovered by the addition of EDTA (with 1:1 EDTA:Zn molar ratio), suggesting the dissociation of the Zn-Aβ(1-40) complex to form monomer-like Aβ(1-40) species and Zn-EDTA complex (Figures 5B and S11B). This is further verified by size profiling using size-exclusion chromatography (SEC). The Aβ(1-40) oligomers prepared using phenol red free F-12 cell-culture media showed an elution profile with three peaks appearing near ∼18-19 mL correspond to oligomers and a single peak near ∼21 mL correspond to monomers (Figure 5C, blue); the SEC Aβ(1-40) monomers incubated for 7 days showed a major population of fibers that eluted at the dead volume (∼ 7 mL), and also a small amount of oligomers (∼15 mL) and monomers (∼21 mL) (Figure 5D, black). cf-E111Q-IDE was observed to elute near ∼8 mL (Figure 5C, red), and Zn-Aβ(1-40) sample depicted monomers, oligomers, and larger-size aggregates (Figure 5C, green). As anticipated, Aβ(1-40) mixed with cf-E111Q-IDE presented monomers, small oligomers, and mostly large aggregates (Figure 5D, blue), which correlates to our TEM and NMR observations. Notably, Aβ(1-40) incubated with zinc+EDTA+cf-E111Q-IDE or EDTA+ cf-E111Q-IDE showed monomers as a major population, some fibers, and no oligomer peak (Figure 5D). Taken together, the SEC, ThT fluorescence, and NMR experiments reveal: (i) zinc-chelation of cf-E111Q-IDE (zinc-free enzyme) does not alter its proteolytic activity on Aβ(1-40), and (ii) removal of zinc from the Zn-Aβ complex by EDTA enables the formation of monomers and low-molecular-weight aggregates of Aβ.

**Figure 5.**
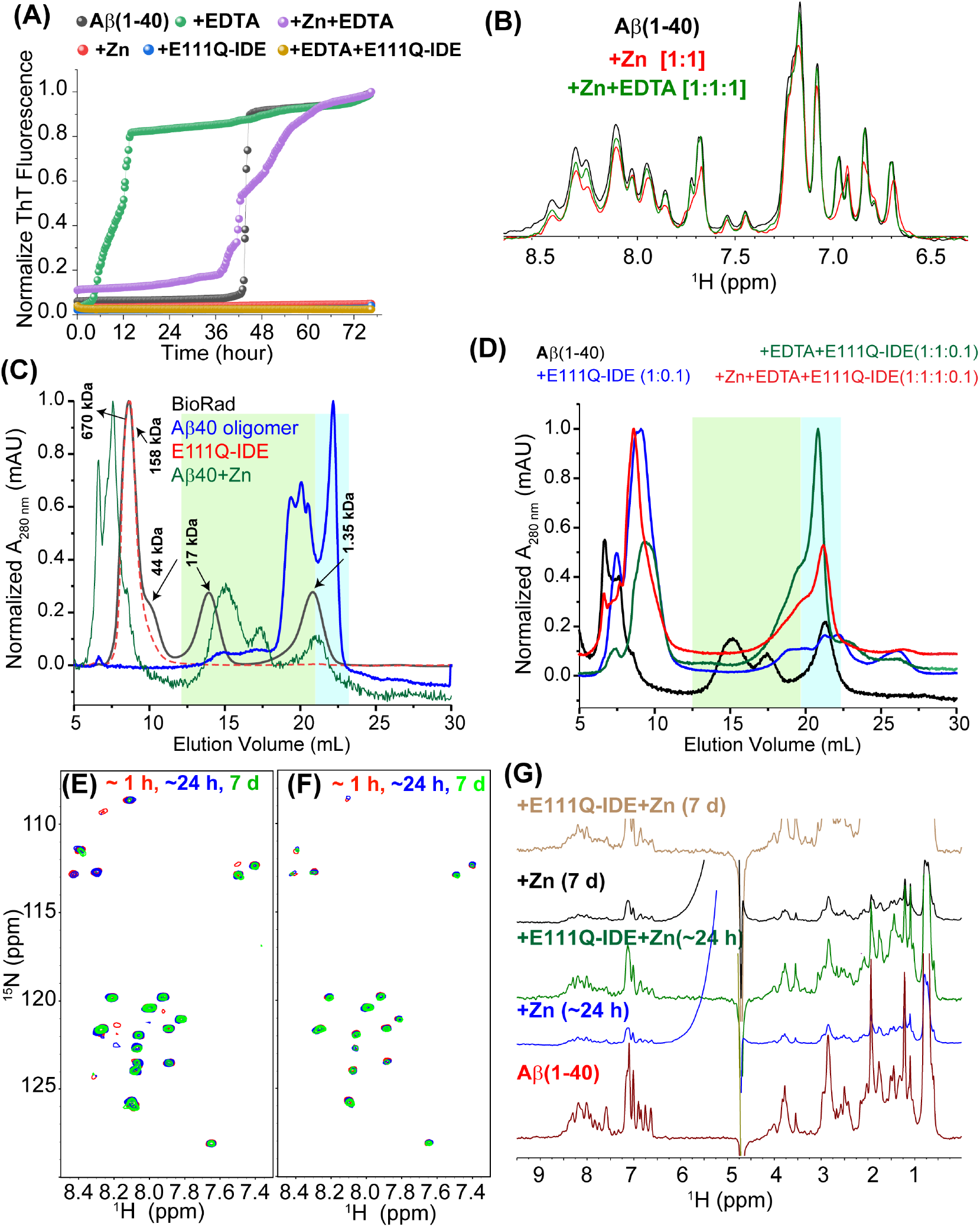
Catalytic effect of E111Q-IDE on Aβ(1-40) in the presence of zinc and EDTA. (A) ThT curve (average from triplicate with error bar is shown in Figure S10) representing the aggregation kinetics of 5 µM Aβ(1-40) in the absence or presence of 5 µM zinc, 5 µM EDTA, and 0.5 µM E111Q-IDE as indicated in colors at 25 °C; for example, the green data points are from the sample of Aβ+EDTA. (B) ^1^H NMR spectra of 25 µM Aβ(1-40) showing a reversible change in signal intensity (see also Figure S11B) upon zinc binding and chelation using EDTA at the indicated stoichiometry. (C-D) Size-profile analysis of 25 µM Aβ(1-40) using size-exclusion chromatography (SEC) in the absence or presence of E111Q-IDE, zinc and EDTA. (C) Aβ(1-40) oligomers, IDE, and BioRad protein ladder are used as reference to characterize Aβ species in the presence of E111Q-IDE, Zn and EDTA. (D) NMR samples used for SOFAST-HMQC measurements (Figures 1, 5, and 6) on day-7 were used for SEC as indicated in colors. The highlighted colors (background vertical bars) correspond to the size distribution indicated in (C). Time-lapse NMR spectra of 25 µM Aβ(1-40) co-incubated with 25 µM zinc in the absence (E) or presence of 2.5 µM E111Q-IDE (F) as indicated in colors. NMR samples were prepared using 10 mM NaPi buffer, pH 7.4 containing 10% D_2_O, and spectra were recorded on a 600 MHz spectrometer at 25 °C. (G) Time-evolved decay in ^1^H NMR signal intensity obtained from 25 µM Aβ(1-40) co-incubated with 25 µM zinc in the absence or presence of 2.5 µM E111Q-IDE as a function of time (see also Figure S11A). E111Q-IDE (sh1o7wn in A, D and G) indicates the cysteine-free E111Q-IDE mutant.

### Zinc bound Aβ(1-40) species is not a substrate for cf-E111Q-IDE

The effect of cf-E111Q-IDE on the Zn-Aβ(1-40) complex was next monitored using 2D SOFAST-HMQC. As shown in Figure 5E, the Zn-Aβ(1-40) complex incubated for ∼1 hour showed a considerable decrease in the signal intensity for the N-terminal residues (A2, E3, Y10, E11, V12, and K16) as compared to the spectrum of Aβ(1-40) observed in the absence of zinc (Figure 1I). This observation is in agreement with zinc-binding and the formation of Zn-Aβ(1-40) complex, as reported previously.[45,46] Time-lapse monitoring further exhibited a decrease in the signal intensity for several residues on day-7 indicating an increase in the size of Zn-Aβ(1-40) aggregates. Growth in the size of Zn-Aβ(1-40) oligomers has also been previously observed with aging time using an electron microscope. [17] To our surprise, the Zn-Aβ(1-40) complex incubated with 1:10 cf-E111Q-IDE:Aβ molar ratios for 7 days showed no substantial change in the NMR resonance pattern. In addition, unlike our observation for Aβ(1-40) proteolysis by cf-E111Q-IDE in the absence of zinc (Figure 1J and L), we did not observe the appearance of new peaks (Figure 2B) or the disappearance of existing peaks until day-7 (Figure 5F). This suggests that the zinc bound Aβ(1-40) species is not a substrate of cf-E111Q-IDE due to its unfavorable size that cannot be accommodated on the catalytic cavity of IDE. The relative changes from day-1 to day-7 were further quantified from ^1^H NMR signal intensities for zinc+Aβ(1-40) and zinc+cf-E111Q-IDE+Aβ(1-40) samples (Figures 5G, S11A and S11B). Integrated intensities of amide peaks (7.5 to 8.5 ppm, Figure 5G) depicted no significant change for both zinc+Aβ(1-40) and zinc+cf-E111Q-IDE+Aβ(1-40) samples. These results rule out the possibility of monomer dissociation, proteolysis, and fibrillation.

### Zinc removal from the Zn-Aβ complex resumes Aβ degradation by cf-E111Q-IDE

Considering the crucial role of zinc in AD pathology progression,[43] our NMR results highlighted a dual function for zinc that has been poorly explored. Zinc binding enhances the toxic activity of Aβ [43,44], and as shown in Figure 5A zinc also impaired IDE’s proteolytic activity. To this extent, we next examined the possibility of reversing the proteolytic activity of IDE through zinc chelation. As discussed in the previous section, EDTA showed an optimal zinc chelating activity at equimolar concentration without affecting the activity of cf-E111Q-IDE. Real-time monitoring using HS-AFM was carried out to probe the efficacy of EDTA to reverse the morphology of Zn-Aβ(1-40) aggregates and generate cf-E111Q-IDE preferred small aggregate or monomer substrates. The zinc bound Aβ(1-40) complex was found to be heterogeneous with size varying from ∼5 nm to several hundreds of nanometer in diameter (Figure 6A-D). Quantitative analysis of particle size from two representative images derived from HS-AFM movie (Figures 6(E and F) and S12) presented a relatively high number of small particles (area 50-200 nm^2^) in sample containing Aβ-Zn-cf-E111Q-IDE. An increase in the particle size to 500-1000 nm^2^ was observed over time (supporting movie SV1) suggesting the assembling of small Zn-Aβ(1-40) particles in the solution (Figure 6G, black bar). On the other hand, EDTA titration substantially reduced the number of small particles (50-200 nm^2^, Figures 6F and S12), as illustrated in Figure 6G (green bars). Unlike the observation of growth in particle size (500-1000 nm^2^), EDTA titration significantly restricted particle fusion (supporting movie SV2) and the formation of large aggregates (Figures 6D and G, blue bar). Taken together, HS-AFM analysis indicated the dissociation of small aggregates (area 50-200 nm^2^) to invisible species, which could be a preferred substrate for IDE. To test if these EDTA induced species are a substrate of cf-E111Q-IDE, we performed time-lapse ^1^H-^15^N SOFAST-HMQC experiments. The appearance of several new peaks in the NMR spectrum of Aβ(1-40)+EDTA+cf-E111Q-IDE at a time-interval of ∼24 hours (Figure 6H) indicates the proteolytic activity of cf-E111Q-IDE on Aβ(1-40) similar to the results obtained in the absence of EDTA (Figures 1J). It is remarkable that the pre-incubated Zn-Aβ(1-40) aggregates, that showed resistance to enzymatic proteolysis (Figure 5F), when titrated with EDTA and cf-E111Q-IDE resulted in the appearance of several new peaks in the NMR spectrum within ∼24 hours (Figures 2B (purple stars) and 6I) indicating Aβ(1-40) fragmentation. This NMR observation correlates to the ThT fluorescence quenching results (Figure 5A, yellow trace) indicating the proteolytic cleavage of zinc-chelated-Aβ(1-40) in the presence of EDTA by cf-E111Q-IDE.

**Figure 6.**
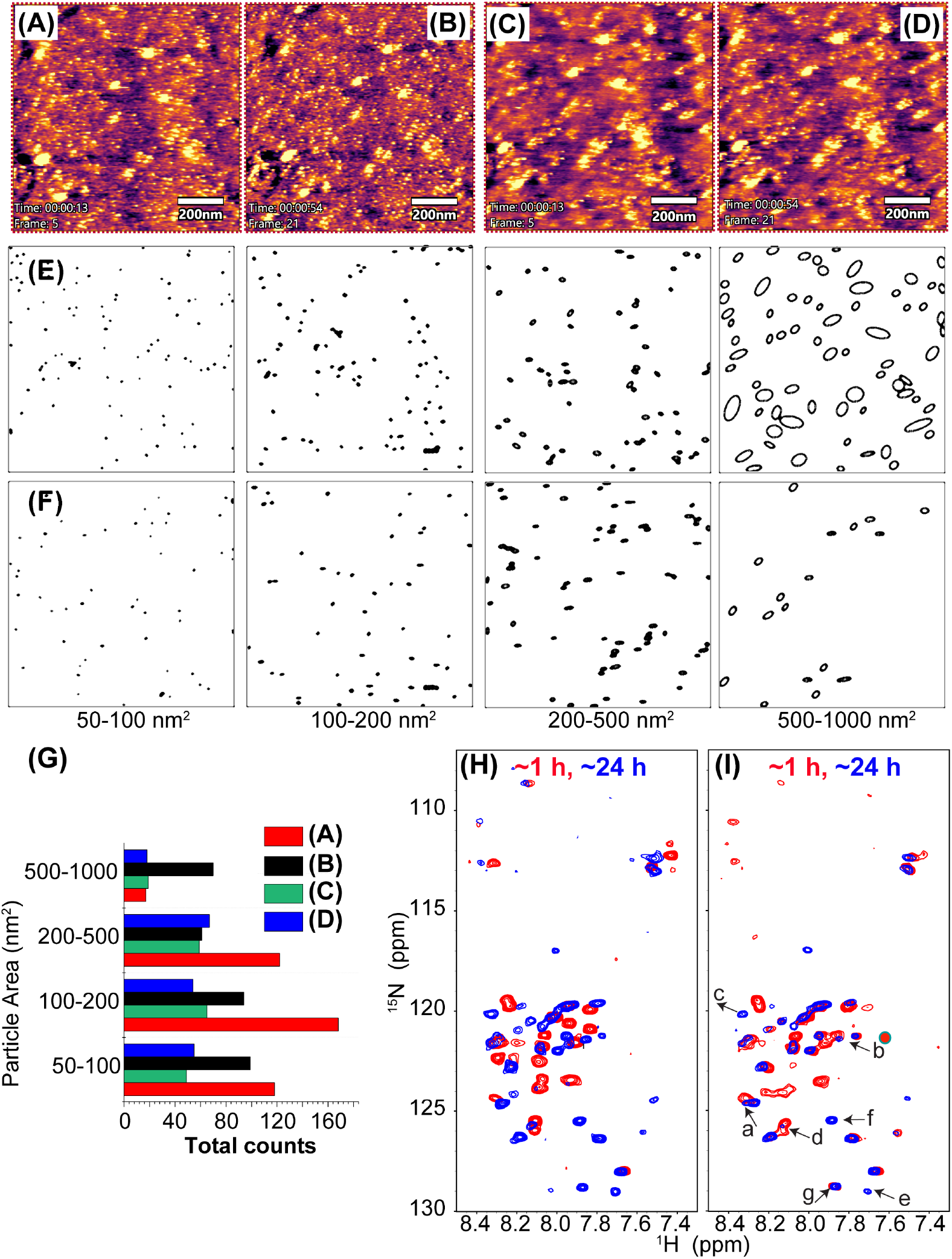
Real-time HS-AFM monitoring of 5 µM zinc-Aβ (1:1) complex incubated with 0.5 µM E111Q-IDE in the absence (A, B) or presence of 5 µM EDTA (C, D). The HS-AFM images are retrieved at different time-points as indicated, are referred to as frame-5 (A or C), and frame-21 (B or D). (E-F) Distribution of particle size derived using the HS-AFM images shown in (B, D) are plotted in ImageJ using the size filters shown on the bottom. The particles shown in (E) corresponds to frame-21 illustrated in (B) that contains Aβ:Zn:E111Q-IDE (1:1:0.1), and particles in (F) corresponds to frame-21 illustrated in (D) that contains Aβ:Zn: EDTA:E111Q-IDE (1:1:1:0.1). The particles retrieved from Frame-5 are shown in Figure S12. (G) Quantitative analysis of the Zn-Aβ-IDE particle size obtained from the HS-AFM images shown in A-D at the indicated colors in the absence or presence of EDTA. SOFAST-HMQC spectra of 25 µM ^15^N labelled Aβ(1-40) co-incubated with 25 µM EDTA and 2.5 µM E111Q-IDE without (H) and with (I) externally added 25 µM zinc recorded at the indicated time intervals. The unassigned newly appeared peaks (labeled as a-g in spectra (I)) are indicated by arrows and the volume ratio (1 vs 24 hr peaks) is plotted in Figure 2B. NMR samples were prepared using 10 mM NaPi buffer, pH 7.4 containing 10% D_2_O, and spectra were recorded on a 600 MHz spectrometer at 25 °C.

## Discussion

Due to the many unsuccessful attempts in drug development to potentially treat Alzheimer’s disease[7] that mainly focused on amyloid inhibition, there is considerable interest in exploring other ways to develop therapeutic strategies. In this context, it is useful to investigate the important roles played by proteolytic enzymes on amyloidogenic protein degradation and clearance. Since the accumulation of Aβ aggregates is found in a majority of AD patients, the correlation between the dysfunction of Aβ peptide degradation/clearance mechanism and Alzheimer’s disease progression was proposed several decades ago.[47,48] The insulin-degrading enzyme is one among several enzymes that control the progression of AD and T2D diseases by degrading Aβ, insulin and amylin peptides in cellular fluids.[38] In view of the diverse cellular localization of IDE, a substantial correlation between IDE and Aβ has previously been demonstrated both in vitro and in vivo. [28,35,49,50] A decrease in cerebral Aβ deposits (> 50%) has been reported in mice that lack IDE.[49] This highlights the important roles played by IDE on Aβ clearance and the pathogenesis of AD. Allosteric modulation of IDE’s catalytic activity is shown to be important to fight against type-2 diabetes and Alzheimer’s disease.[33] To selectively enhance the degradation of Aβ by IDE, a key strategy employed was to allosterically modulate IDE’s activity by using compounds which were further validated through in-vitro mutational studies.[32] IDE mutants such as E111Q were shown to be inactive against insulin degradation indicating their importance for better understanding the influence of IDE on the role of insulin in type-2 diabetes. Although the development of non-catalytic IDE mutants is important, their effect on the degradation and clearance of Aβ need to be examined. Specifically, the design of IDE-mutants to validate allosteric modulation of IDE should make sure that they are capable of degrading Aβ like the wild-type IDE. To address this point, in this study we undertook an investigation to evaluate the effect of one such IDE mutant, cf-E111Q-IDE, on the amyloid aggregation of Aβ. A remarkable finding in this study is that the cf-E111Q-IDE mutant enzymatically degrades Aβ peptides.

A previous study proposed chaperone activity of E111Q-IDE following the observation of its effect on delaying Aβ aggregation in-vitro.[35] In this study, we investigated the effects of both WT-IDE and the mutant cf-E111Q-IDE using a variety of biophysical approaches. The ThT fluorescence results show a significant delay in Aβ aggregation (Figures S2 and S3) by both wild-type-IDE and the mutant cf-E111Q-IDE which is in agreement with previous studies.[28,35] Unlike the WT-IDE, the cf-E111Q-IDE mutant exhibited a comparatively smaller effect on Aβ(1-40) fibrillation at substoichiometric concentration, 1:100 molar ratio of enzyme:peptide (Figure S2). According to previous reports [23, 39], the reduced activity of the mutant observed at a substoichiometric concentration could be due to cysteine-deletion and E111Q mutation. However, at a higher concentration 1:10 enzyme:peptide molar ratio, the cf-E111Q-IDE mutant showed no aggregation of Aβ(1-40) in ThT experiments similar to that observed for the WT-IDE (Figure S3) which is because of the degradation of the peptide as confirmed by LC-MS (Figure 1) and not due to chaperone activity as proposed by the previous study.[35] To further examine the non-chaperone activity of the cf-E111Q-IDE, a series of experiments that include LC-MS, DOSY NMR, SOFAST-HMQC and TEM were carried out. LC-MS revealed the presence of short sized Aβ fragments in the presence of cf-E111Q-IDE that are similar to what we observed in the presence of WT-IDE (Figure 1(A and B)). Real-time diffusion NMR experiments identified the presence of small Aβ fragments whose diffusion rates similar to that for tri/tetra peptide molecules (used as reference) in the presence of wild-type or E111Q mutant IDE (Figures 2(C-F) and S7). The 2D SOFAST-HMQC NMR spectra revealed the appearance of additional narrow resonances appearing from peptide fragments as evidenced from DOSY and LC-MS data. These fragmented peptides were found to aggregate over time as they showed progressive line-broadenings, which is in agreement with a previous NMR study on the effect of WT-IDE on Aβ.[28] TEM images identified a morphologically similar species varying in size ∼20-30 nm from both wild-type and mutant IDE samples (Figure 1F and G). In sum, these results clearly confirm the non-chaperone activities for both wild-type and mutant IDEs.

Spherically shaped oligomeric species formed by the full-length Aβ have been shown to be toxic.[51] Whereas the aggregates of Aβ fragments, cleaved by the WT-IDE or the cf-E111Q-IDE mutant, have been reported to be non-toxic [28], but possess spherical morphology as seen from the TEM images reported here (Figure 1E-H). Therefore, it is unclear why these morphologically similar oligomers formed by the full-length and fragments exhibit pathologically different phenotypes. The toxicity of Aβ oligomers in the self-assembled and hetero-assembled states has been proposed to be influenced by their exposed hydrophobic surface that stimulate membrane binding. [42,52,53] Similarly, polymorphic Aβ fibrils,[54] Aβ oligomers trapped in the membrane, [55] and metal-coordinated oligomers [10,44] have been shown to exhibit variable neurotoxicity. To understand why the morphologically similar oligomers exhibit difference in toxicity, we carried out molecular dynamics simulations on the fragmented and full-length Aβ peptides. Our results indicate that there exists a unique common feature between the full-length Aβ(1-40) aggregates and the fragmented Aβ aggregates (Figure 4). Aggregates of the full-length Aβ(1-40) or long-fragments (Figure 3A) are characterized by a decrease in SASA value and exposure of central hydrophobic residues 21-28. Whereas the short N- or C-terminal fragments like Aβ(1-12), Aβ(1-13), Aβ(1-14), Aβ(34-40), Aβ(35-40) and Aβ(36-40) possess a large SASA value. Notably, the N-terminal truncated Aβ(13-40) aggregates showed a buried central hydrophobic region in contrast to the full-length Aβ(1-40). This indicates that the cleavage of the highly charged N-terminal fragment Aβ(1-13) by IDE generates a fragment that opposes the exposure of central hydrophobic residues which are proposed to influence Aβ toxicity.[51] Further, the atomic-resolution structure of Aβ peptide assembly revealed large-size aggregates for the full-length Aβ (Figure 4A), whereas Aβ-fragments are found as monomers or low-order aggregates in the solution. The distinguished molecular feature of Aβ/Aβ-fragment self-assembly obtained from the atomic-resolution simulation are correlating well with the diffusion NMR results (Figure 2E and F) that detected short Aβ-fragments in solution. These simulation results provide a mechanistic detail highlighting a considerable difference in the solvent-accessibility between the spherical Aβ/Aβ-fragment oligomers, which could possibly be associated with their opposite neurotoxic activities.

The toxicity of Aβ peptide is highly modulated by the presence of metals that include copper, zinc, iron, aluminum, etc. and have been extensively studied in the past few decades.[10] However, less has been explored how metals influence the activity of the enzymatic machinery that recycles the soluble amyloidogenic peptides. Our *in-vitro* results show that zinc impairs the proteolytic activity of cf-E111Q-IDE against Aβ. Also, the activity of cf-E111Q-IDE is impaired by zinc ions that are shown to be present up to millimolar concentration in a cellular milieu.[56] Therefore, our data indicate that zinc induces the formation of stable zinc bound Aβ oligomers (Figure 5E and F) which cannot fit the catalytic chamber of cf-E111Q-IDE. Thus, our results demonstrate how the aggregation pathway could be redirected to resume the proteolytic activity of IDE (Figure 7). The impairment of the enzymatic degradation is proposed to be stimulated by an increase in the Aβ hydrophobicity upon zinc binding in familial AD.[24] Therefore, the therapeutic intervention of the formation of metal-stabilized Aβ oligomers could be a potential alternative strategy to control the disease progression. Although a direct correlation between aging Aβ species and enzymatic degradation has been established,[25] development of strategic zinc-chelating small-molecule targeting the IDE-Zn-Aβ complex is not well investigated. This in-vitro study demonstrated the activity of the cf-E111Q-IDE on Aβ proteolysis in real-time using high-speed AFM and NMR (Figure 6). As illustrated in Figure 7, EDTA, a known zinc chelator, has been shown to disassemble zinc-bound Aβ aggregates as evidenced from HS-AFM and NMR (Figures 5B and 6D). The small and large size Zn-Aβ aggregates in the presence of EDTA disintegrate to form monomers or low-order aggregates (Figure 6(C-D)) that are identified to be degraded by cf-E111Q-IDE in NMR (Figure 6I). On the basis of the findings presented in this study, we summarize that the formation of the zinc-Aβ complex but not E111Q-mutation impairs the enzymatic activity of cf-E111Q-IDE (Figure 7). These findings highlight that the cf-E111Q-IDE mutant could be a potential therapeutic alternative to treat AD and T2D subjected to the development of potential zinc-chelating drug candidates. Additionally, we expect that the reported findings to have high impacts in the field as an increase in the risk of T2D in AD [57] or vice-versa and cross-seeding among various amyloidogenic proteins, including Aβ and amylin, remarkably highlight the pathological correlation between AD and T2D.[58]

**Figure 7.**
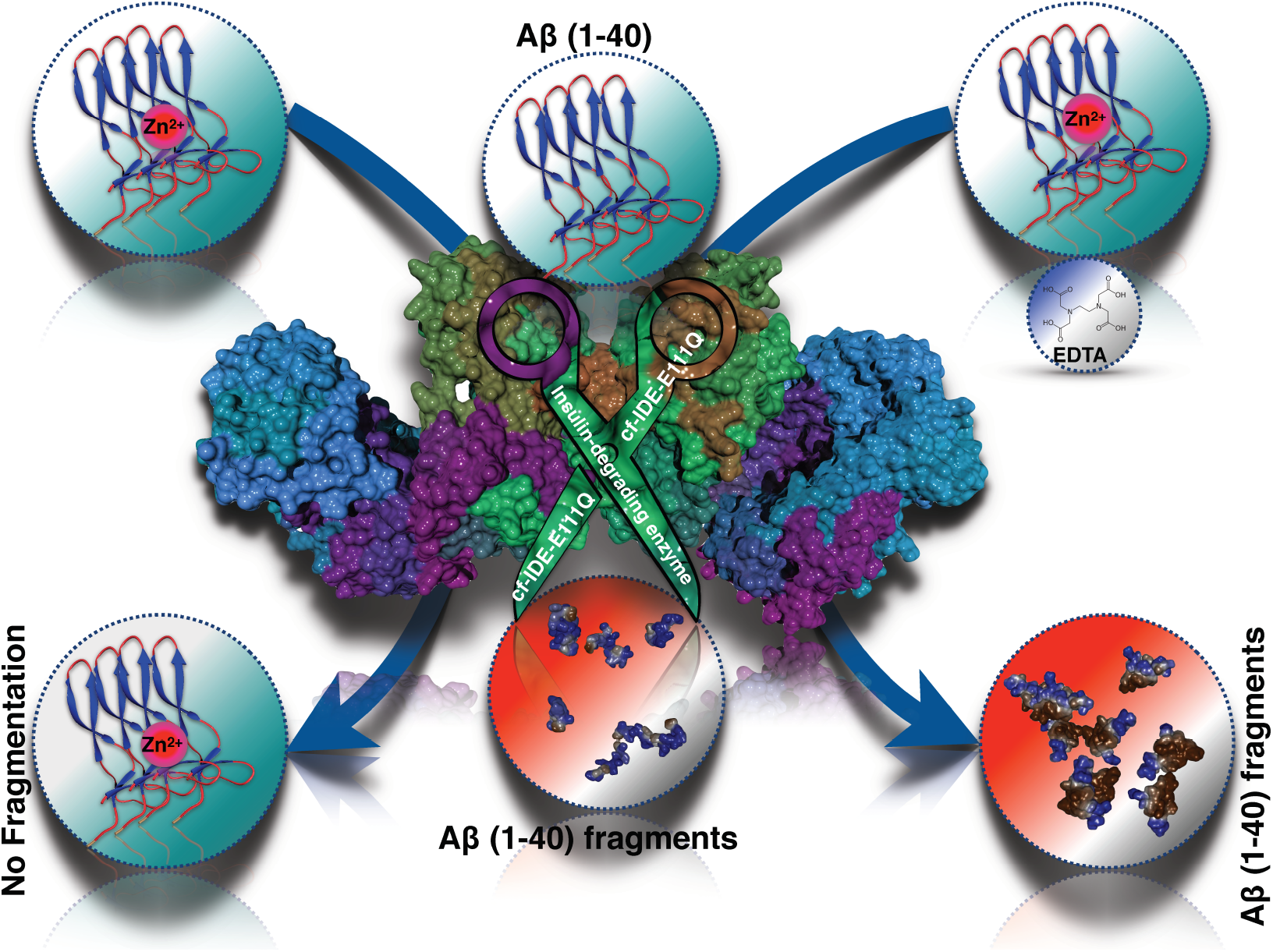
A schematic showing how zinc impairs IDE’s catalytic activity by forming Zn-Aβ complex that does not fit to the IDE’s ≈35 Å active site. Zinc removal from the Zn-Aβ complex by EDTA enables the degradation of Aβ by cf - E111Q-IDE. The degraded Aβ fragments self-assemble to form aggregates of size > 20 nm or do not aggregate in solution depending on the cleavage site.

In conclusion, we have demonstrated the proteolytic activities of wild-type and cysteine-free E111Q mutant IDEs on Aβ(1-40). Both IDEs were found to be catalytically active and able to cleave Aβ(1-40) monomers to short fragments. The cleaved Aβ fragments are shown to self-assemble to form morphologically similar globular structures of size ∼20-30 nm that have been previously tested to be nontoxic.[28] Atomic simulation results revealed small SASA values for the full-length Aβ spherical aggregates as compared to the aggregates formed by the Aβ-fragments, which explained their distinct toxic phenotypes at a molecular level. Another key finding of this study is the ability of zinc to generate proteolytically resistant Aβ(1-40) species that are pathologically relevant (Figure 7). Zinc binding to Aβ was found to form off-pathway aggregates that grow in size and do not fit to the catalytic cavity (≈3.5 nm) of cf-E111Q-IDE (Figure 7). Zinc removal from the Zn-Aβ complex was shown to resume the proteolytic activity of cf-E111Q-IDE. Taken together, these findings highlight the key role of zinc in AD in incapacitating the enzymatic degradation of soluble Aβ monomers. The results reported in this study urge for the development of potent therapeutic zinc chelating agents to resume the enzymatic degradation of soluble Aβ and prevention of AD progression.

## Materials and methods

### Expression and purification of Aβ(1-40) and IDE Proteins

Full-length unlabeled and uniformly ^15^N isotope labeled Aβ(1-40) (MDAEFRHDSGYEVHHQKLVFFAEDVGSNKGAIIGLMVGGVV-NH_2_) with an extra methionine residue at the N-terminus ((known to have no effect on the Aβ(1-40) aggregation kinetics)) were recombinantly expressed in *E. coli* BL21 (DE3). The expression for Aβ(1-40) was adopted from a previously described method [59] and purified by loading the samples to an ^ECO^PLUS HPLC column packed with the reversed-phase separation material. The purification protocol described elsewhere was used in this study.[59] The fractions containing the purified Aβ(1-40) were verified using SDS-PAGE, and peptide samples were lyophilized. The lyophilized Aβ(1-40) powder was next treated with 5% (v/v) NH_4_OH and lyophilized at a concentration of 0.1 mg/mL for ∼48 hours. The peptide was dissolved in 10 mM sodium phosphate (NaPi) buffer, pH 7.4 following a short vortex mixing, ∼15 s sonication and centrifugation at 14,000 × g for 15 min at 4 °C. Oligomers of Aβ(1-40) were prepared by freshly dissolving the peptide powder in 50 µL of 10 mM NaPi, pH 7.4 (monomers) followed by dilution in phenol red free F-12 cell-culture media (950 µL). The peptide sample was incubated at 4°C for 24 hours in dark prior to size-exclusion chromatography. All other chemicals were purchased from Sigma Aldrich, with a purity greater than 95% and used without further purification.

Wild-type and the cf-E111Q-IDE mutant proteins were expressed in *E. coli* BL21 (DE3) cells (at 25 °C and 20 h, 0.5 mM IPTG induction using T7 medium). Recombinant IDE proteins were purified by Ni-NTA, source-Q, and Superdex 200 columns as described in the literature. [60] The IDE enzyme was re-suspended in 10 mM NaPi buffer by exchanging the buffer using a 30-kDa amicon filter. Unless and otherwise mentioned, all the experiments were carried out using 10 mM NaPi, pH 7.4. The enzymatic activity of purified IDE (2.5 nM) was accessed by testing the hydrolysis of 2.5 µM Substrate V (7-methoxycoumarin-4-yl-acetyl-RPPGFSAFK-2,4-dinitrophenyl, R&D Systems), a previously known fluorogenic substrate of IDE. The activity of 2.5 nM WT-IDE or cf-E111Q-IDE co-incubated with 20 mM EDTA was also tested on the hydrolysis of substrate V. Specifically, the reaction was carried out in the presence of 5 nM IDE in 100 μl 20 mM HEPES, pH 8.0, 150 mM NaCl with/without 20 mM EDTA mixed with 5 μM substrate V in 100 μl 20 mM HEPES, pH 8.0, 150 mM NaCl in a final volume of 200 μl at 37 °C. The hydrolysis of substrate V was measured every 30 seconds for 10 minutes using a fluorometer (Synergy Neo HST Plate Reader) with excitation and emission wavelengths set at 320 nm and 405 nm, respectively. The catalytic activity of IDE in the absence and presence of equimolar EDTA (0.5µM) incubated overnight was also tested on 5 µM Substrate-V as a function of time.

### ThT fluorescence assay

The amyloid fibrillation kinetics of 5 µM Aβ(1-40) was monitored using Thioflavin T (ThT) dye based fluorescence assay at 25 °C in the presence and absence of wild-type or IDE mutant. 2.5 or 5 µM of Aβ(1-40) dissolved in 10 mM NaPi buffer containing 10 µM ThT was used for the fluorescence measurements. Enzyme:peptide stoichiometries of 1:10, 1:250, and 1:1000 were used to test the catalytic activity of IDE on Aβ(1-40). A 96-well polystyrene plate (Fisher) with a sample volume of 100 μL/well was used for the ThT fluorescence measurements under medium shaking conditions. ThT measurements were also performed for the IDE enzymes co-incubated with 5 µM of EDTA overnight following the addition to 5 µM of Aβ(1-40) in the ThT plate. The fluorescence data points were collected for every 5-min interval using a microplate reader (Biotek Synergy 2) with an excitation and emission wavelengths of 440 and 485 nm, respectively. The plate containing 1:10 IDE:Aβ sample was stored at room temperature, and fluorescence emission was recorded for 30 minutes until day-12 after 60 hours of continuous (every 5 minutes) data collection at 25 °C.

### Transmission electron microscopy (TEM)

5 µM of Aβ(1-40) dissolved in 10 mM NaPi buffer was mixed with 0.5 µM of WT-IDE, cf-E111Q-IDE, or the cf-E111Q-IDE mutant co-incubated with 5 µM of EDTA (overnight at 4 °C). The sample mixture was incubated overnight (∼12 hrs) at room temperature prior to TEM imaging at 25 °C. The TEM sample preparation was followed as described elsewhere. [52] Briefly, 20 μL of sample mixture was added to a collodion-coated copper grid and incubated for ∼5 minutes followed by rinsing with double deionized water. The grid was next stained with 5 μL of 2% (w/v) uranyl acetate, incubated for ∼2 minutes, rinsed with double deionized water followed by overnight drying under vacuum. TEM images were taken using a HITACHI H-7650 microscope (Hitachi, Tokyo, Japan).

### LC-MS

HPLC mass-spectroscopy (LC-MS) was (Agilent) performed with positive ion electrospray ionization mode to identify short Aβ fragments cleaved by IDE. 50 µM of Aβ(1-40) dissolved in 10 mM NaPi buffer was incubated with WT-IDE (5 µM), cf-E111Q-IDE (5 µM) or cf-E111Q-IDE-EDTA (5 µM enzyme, 50 µM EDTA) for ∼12 hours at room temperature. The sample mixture was next filtered using a 10 kDa amicon centrifuge filter. The washed flow through was collected. µ-C18 ZipTip was used to prepare the LC-MS sample following the user manual. The ZipTip was equilibrated with buffer containing 100% acetonitrile (3 times), next with 0.1% formic acid (3 times) followed by sample (washed flow through sample) loading. The loaded sample was next washed with buffer containing 0.1% formic acid (6 times) and eluted using buffer containing a mixture of 60% acetonitrile and 0.1% formic acid.

### NMR experiments

Proton and diffusion ordered spectroscopy (DOSY) NMR experiments were recorded on a 500 MHz Bruker NMR spectrometer equipped with a z-axis gradient triple -resonance (TXO) probe at 25 °C. 2D heteronuclear (^15^N/^1^H) SOFAST-HMQC [61] NMR experiments were performed on a 600 MHz Bruker using a z-axis gradient cryogenic probe. All NMR experiments were performed using 25 µM of unlabeled or labeled Aβ(1-40) dissolved in 10 mM NaPi buffer containing 10% D_2_O at 25°C in the absence and presence of equimolar Zn, EDTA, and 5 or 2.5 or 0.25 µM cf-E111Q-IDE. Proton NMR spectra were recorded with 512 scans and a 2 s recycle delay. 2D SOFAST-HMQC NMR experiments were acquired with 32 scans and 256 t1 increments with a recycle delay of 0.2 s using a Shigemi tube (volume ∼250 µL). The DOSY NMR spectra were recorded using stimulated-echo with bipolar gradient pulses for diffusion and using a gradient strength incrementing from 2 to 98%, 16 gradient strength increments, 36,000 time-domain data points in the t_2_ dimension, 3 s recycle delay, and 100 ms diffusion delay. DOSY NMR spectra of reduced L-Glutathione (0.5 mg/mL) and (Ala)_4_ peptide (0.5 mg/mL) dissolved in 10 mM NaPi, Ph 7.4 containing 10% D_2_O were obtained using the same parameters as used for Aβ or Aβ+cf-E111Q-IDE. All NMR spectra were processed using Bruker Topspin 3.5. NMR intensity plot analysis was performed using MestReNova 12.0.4 (Mestrelab Research S.L.), and DOSY diffusion plot analysis using Bruker Topspin Dynamics center. 2D heteronuclear NMR spectra were analyzed using Sparky (https://www.cgl.ucsf.edu/home/sparky/).

### Molecular dynamics (MD) simulation

All-atom MD simulation of full-length and Aβ fragments were performed in Gromacs 2019.6v[62] using the Swedish National Infrastructure for Computing (SNIC) computing. An extended conformation of Aβ(1-40) obtained from 2 µs MD simulation of the partially folded NMR structure (PDB: 2LFM) was considered as an initial structure (Figure S8). A cubic box with varying sizes containing 9 copies of Aβ/Aβ-fragments was considered for MD simulation using CHARMM36-ff. [63] The MD parameters and protocol were adopted from our previous studies, as described elsewhere. [52] A total of 21 MD systems containing variable-sized Aβ fragments were generated, and 500 ns production MD run for the individual system was carried out at 310 K. The MD trajectory was analyzed using in-house Gromacs commands. 3D structure visualization and depictions were carried out using Chimera v1.14.39. Statistical analysis and plots are generated using Origin 2019 (9.6.0.172).

### Size-exclusion Chromatography (SEC)

The size distribution analysis of Aβ(1-40), cf-E111Q-IDE, and Aβ+cf-E111Q-IDE in the absence and presence of equimolar Zn and EDTA were profiled using SEC. The NMR samples used for 2D SOFAST-HMQC (∼250 μL) measurements were diluted to a total volume of 1 mL using 10 mM NaPi buffer, pH 7.4. The diluted sample was then injected onto a Superdex 75 (10/300GL) column (GE Healthcare) and eluted at a flow rate of 1 mL/min. The size-exclusion chromatography profiles were analyzed and plotted using the Origin program. Bio-Rad’s gel filtration standard mixture (Figure 5C, black) containing thyroglobulin, γ-globulin, ovalbumin, myoglobin, and vitamin B12 (MW 1.35 to 670 kDa) was used as a reference for molecular size comparison.

### High-speed Atomic Force Microscopy (HS-AFM)

The growth of Aβ(1-40) species complexed with zinc was monitored on real-time using the Asylum Research Cypher AFM. HS-AFM instrument operated on tapping mode is equipped with an Olympus BL-AC10DS-A2 microcantilever of spring constant *k*=0.1 Nm^-1^ and resonant frequency f=∼1.5 MHz in water. 5 µM of freshly dissolved Aβ(1-40) monomers dissolved in 10 mM NaPi buffer was mixed with 5 µM of zinc and 0.5 µM of cf-E111Q-IDE and incubated for ∼15 minutes at room temperature. 20 µL of the sample mixture was next added to pre-mounted V4 mica and incubated for ∼10 minutes prior to HS-AFM video rate scanning. Next, 5 µM of EDTA dissolved in 10 mM NaPi was injected to the mica surface using the Cypher VRS syringe following HS-AFM imaging. The HS-AFM videos were analyzed using AR16.10.211 software supported with Igor 6.37 and ImageJ (NIH).

## Author Contributions

Conceptualization: B.R.S., A.R.; Experimental and computational analyses: B.R.S., P.K.P., W.L.; interpretation of results: B.R.S., A.R.; manuscript writing: B.R.S., P.K.P; A.R.; W.J.T, R.A., Y.K.M.; supervision: A.R, R.A; funding acquisition: A.R.

## Acknowledgements

This study was supported by funds from NIH (AG048934 to A.R.). The authors acknowledge the financial support from the Swedish Research Council, (VR grant no. 2016–06014) for providing computational resources, and Asylum Research for the HS-AMF support. We thank Dr. Andrea Stoddard for help with Aβ peptide expression and purification.

