## Supporting information for "Degradation of Alzheimer’s amyloid-β by a catalytically inactive insulin-degrading enzyme"

**Table of Content**:

1. Supporting figures (Figures S1-S14)
2. Supporting Videos (SV1 and SV2)

### 3. Supporting figures

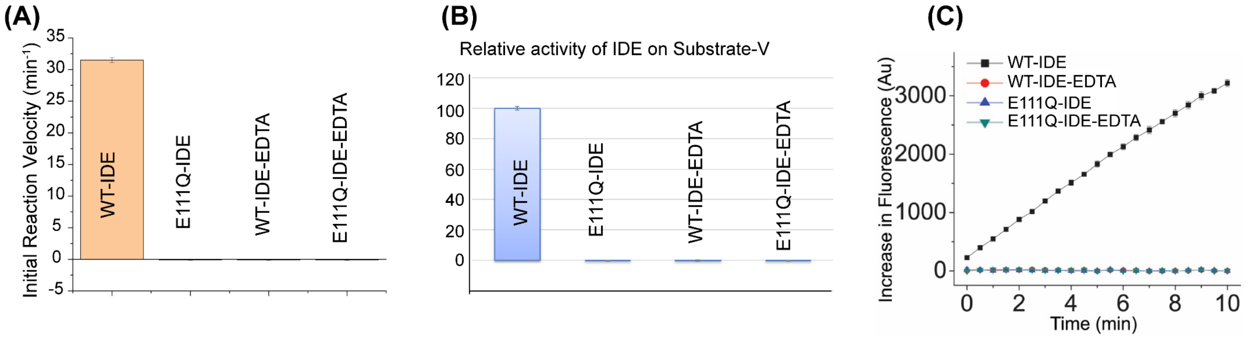


**Figure S1.** Initial reaction velocity (A) and relative activity (B) of 2.5 nM wild-type IDE (labeled as WT-IDE) and cysteine-free-E111Q-IDE mutant (labeled as E111Q-IDE) co-incubated with and without 20 mM EDTA (labeled as WT-IDE-EDTA and E111Q-IDE-EDTA) on 2.5 μM substrate V (a fluorogenic peptide substrate of IDE) measured at 37 °C. (C) Plot showing the change in fluorescence as a function of time in the presence and absence of EDTA.

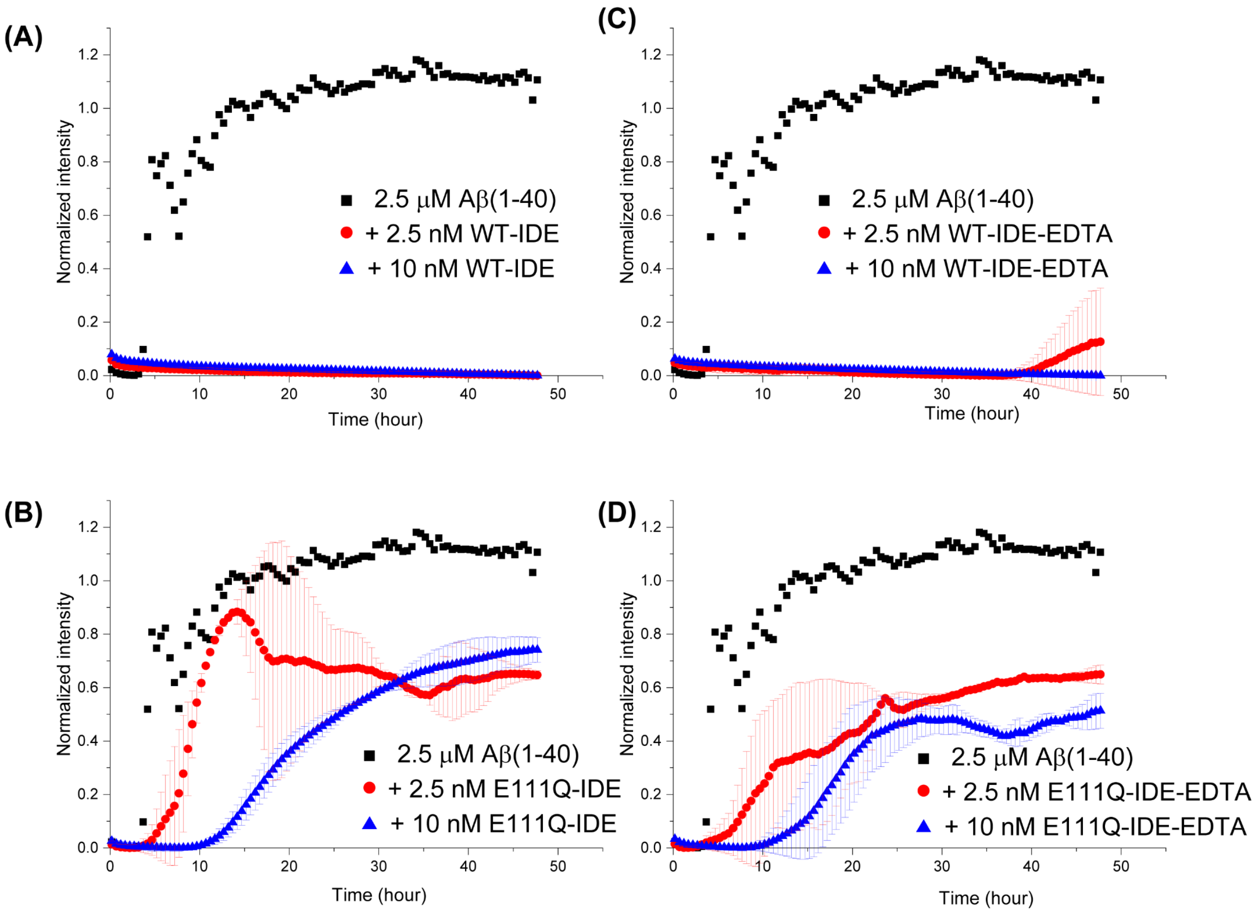


**Figure S2.** Aggregation kinetics of 2.5 μM Aβ(1-40) dissolved in 10 mM NaPi buffer, pH 7.4 in the absence (black) and presence of a variable concentration of wild-type (labeled as WT-IDE) or cysteine-free-mutant (labeled as E111Q-IDE) IDE co-incubated without (A, B) and with (C, D) 2.5 μM of EDTA as indicated. The fluorescence measurements were carried out in triplicate at 25 °C under medium shaking conditions.

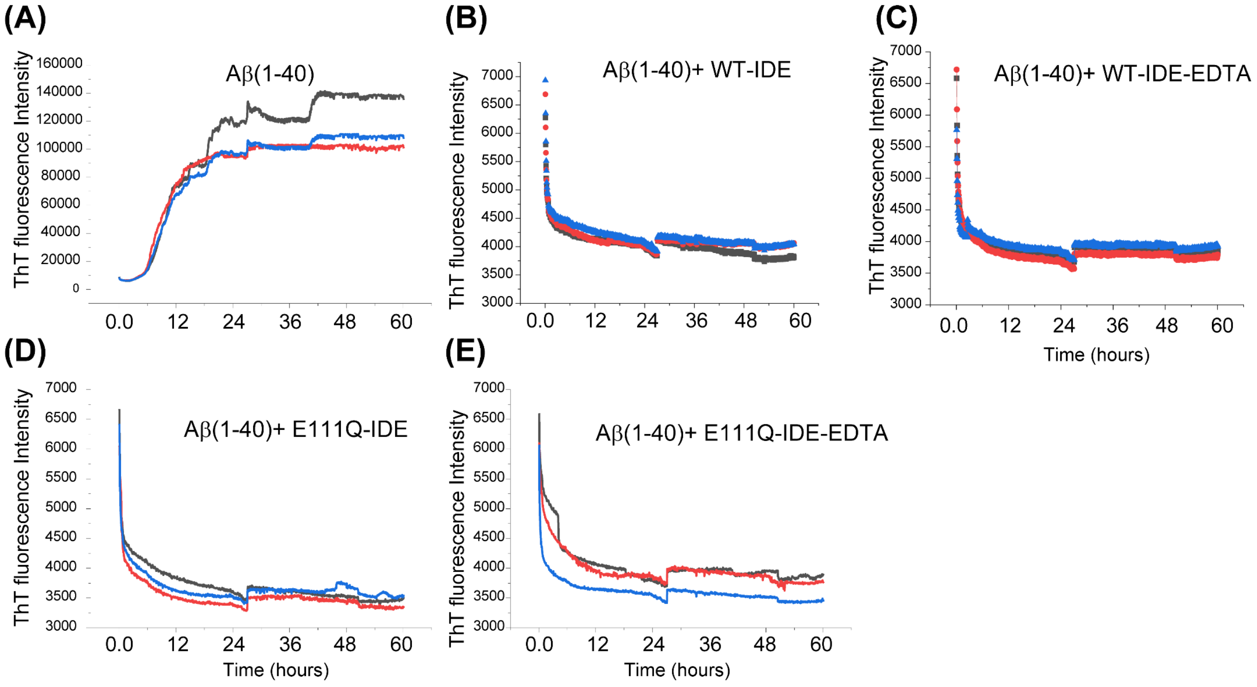


**Figure S3.** ThT fluorescence assay of 5 μM Aβ(1-40) fibrillation. Aggregation kinetics of 5 μM Aβ(1-40) dissolved in 10 mM NaPi buffer, pH 7.4 in the absence (A) and presence of 0.5 μM wild-type IDE co-incubated without (B) and with (C) 5 μM of EDTA. Results obtained in the presence of the cysteine-free E111Q-IDE mutant are shown without (D) and with (E) 5 μM of EDTA. All fluorescence measurements were recorded in triplicate as shown in different colors. The fluorescence measurements were carried out at 25 °C under medium shaking conditions. Data points collected after 60 hours (until day-12) are not shown.

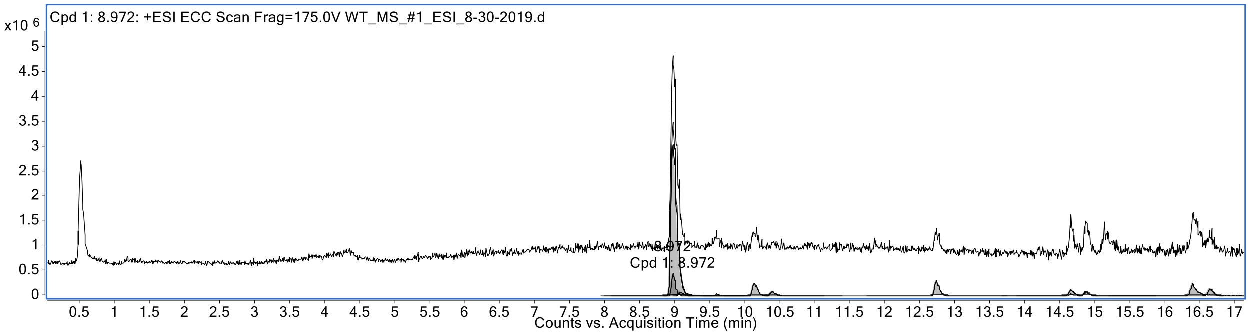
Cpd 1: 8.972: +ESI ECC Scan Frag=175.0V WT_MS_#1_ESI_8-30-2019.d
x10 6
8.972
Cpd 1: 8.972
Counts vs. Acquisition Time (min)

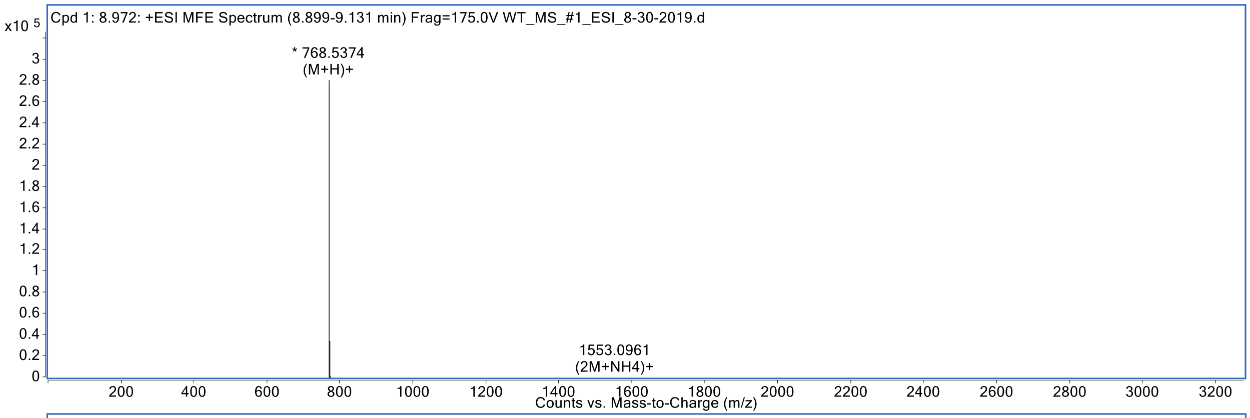
Cpd 1: 8.972: +ESI MFE Spectrum (8.899-9.131 min) Frag=175.0V WT_MS_#1_ESI_8-30-2019.d
x10 5
* 768.5374
(M+H)+
1553.0961
(2M+NH4)+
Counts vs. Mass-to-Charge (m/z)

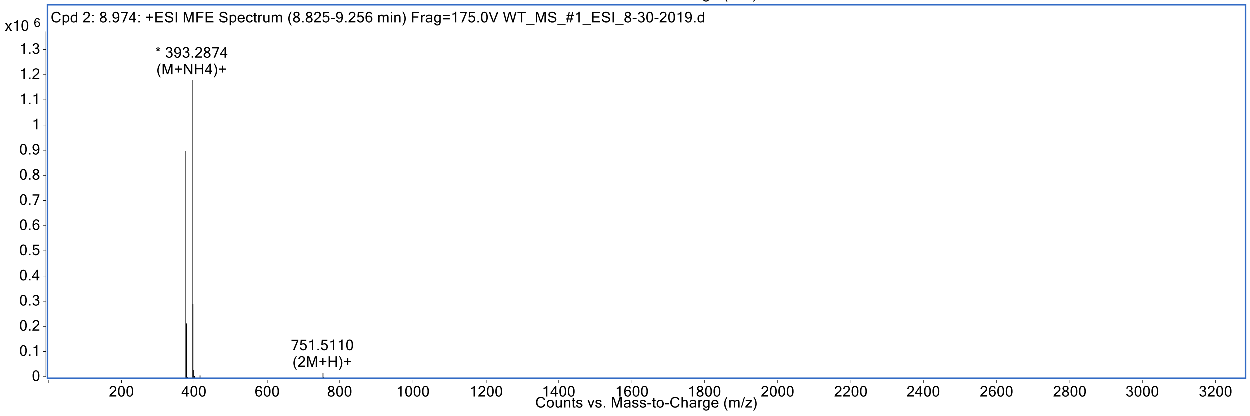
Cpd 2: 8.974: +ESI MFE Spectrum (8.825-9.256 min) Frag=175.0V WT_MS_#1_ESI_8-30-2019.d
x10 6
* 393.2874
(M+NH4)+
751.5110
(2M+H)+
Counts vs. Mass-to-Charge (m/z)

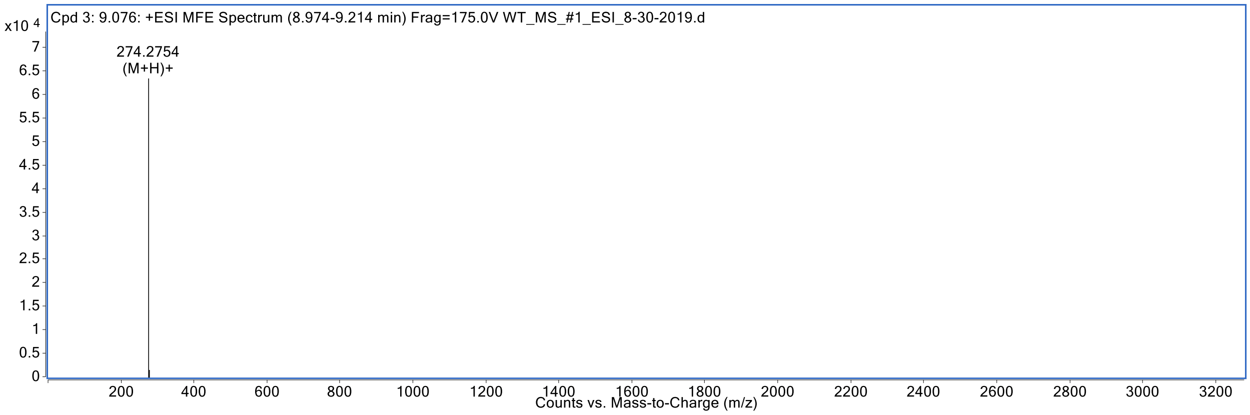
Cpd 3: 9.076: +ESI MFE Spectrum (8.974-9.214 min) Frag=175.0V WT_MS_#1_ESI_8-30-2019.d
x10 4
274.2754
(M+H)+
Counts vs. Mass-to-Charge (m/z)

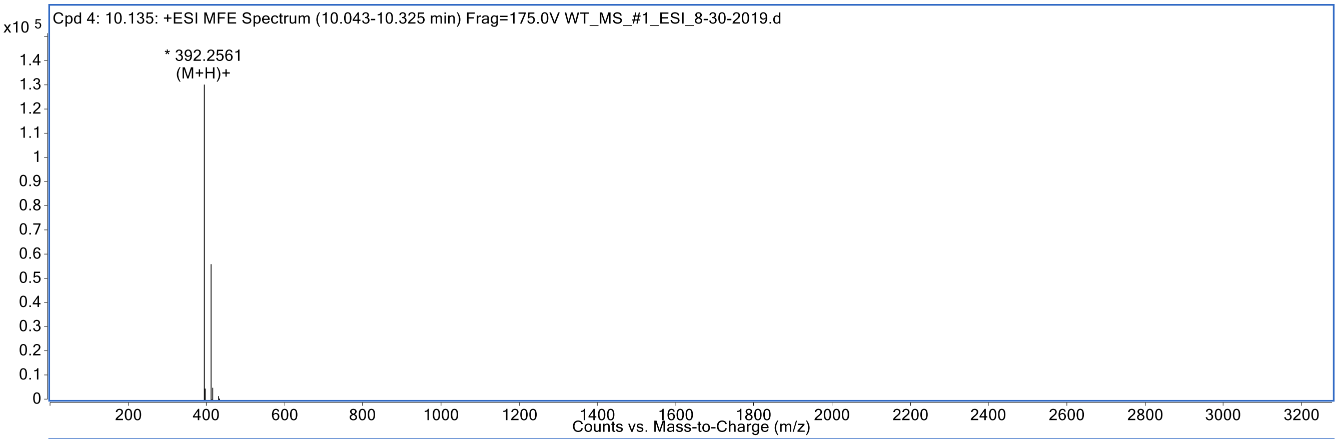
Cpd 4: 10.135: +ESI MFE Spectrum (10.043-10.325 min) Frag=175.0V WT_MS_#1_ESI_8-30-2019.d
x10 5
* 392.2561
(M+H)+
Counts vs. Mass-to-Charge (m/z)

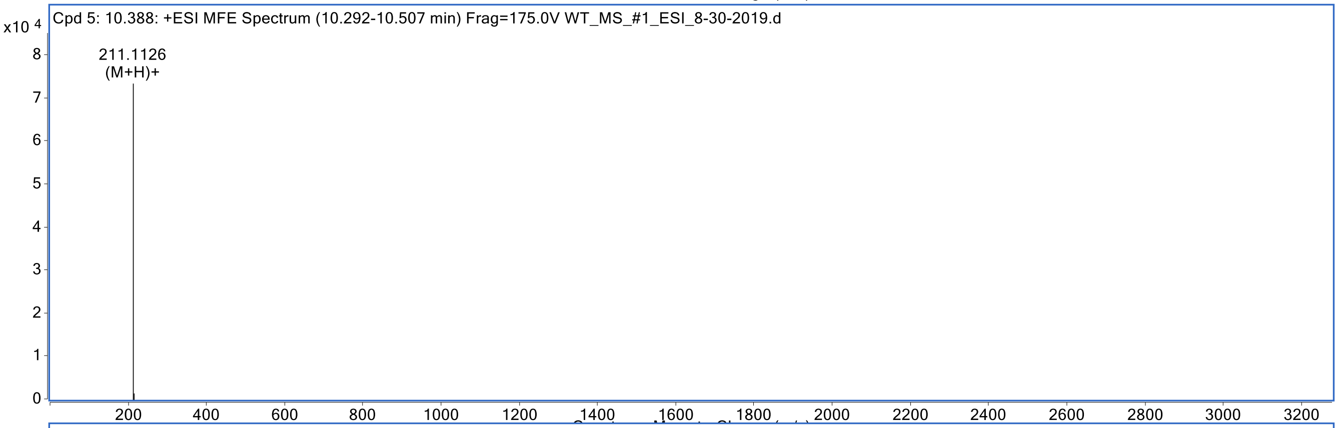
Cpd 5: 10.388: +ESI MFE Spectrum (10.292-10.507 min) Frag=175.0V WT_MS_#1_ESI_8-30-2019.d
x10 4
211.1126
(M+H)+

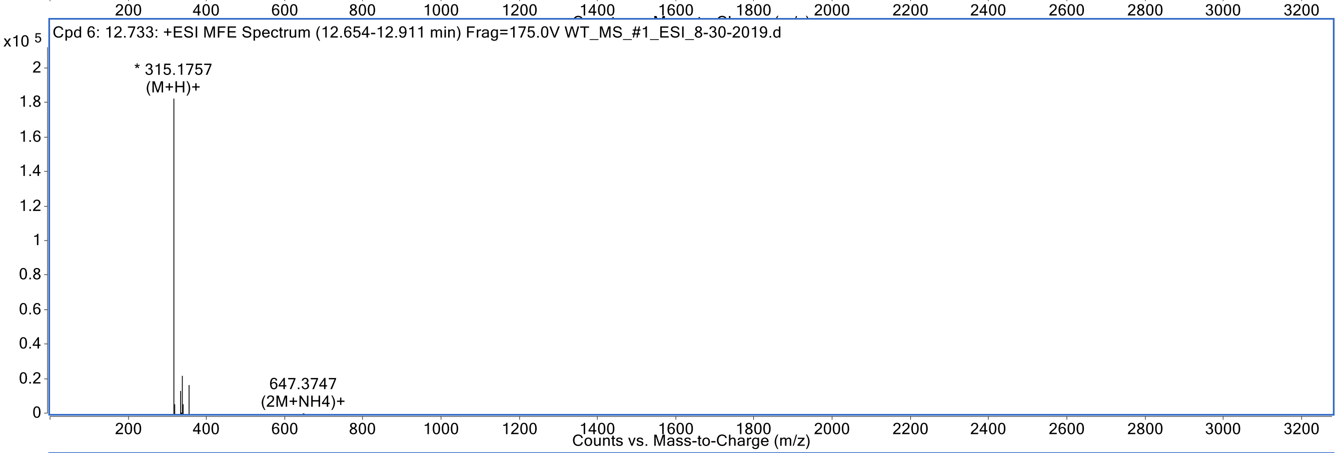
Cpd 6: 12.733: +ESI MFE Spectrum (12.654-12.911 min) Frag=175.0V WT_MS_#1_ESI_8-30-2019.d
x10 5
* 315.1757
(M+H)+
647.3747
(2M+NH4)+
Counts vs. Mass-to-Charge (m/z)

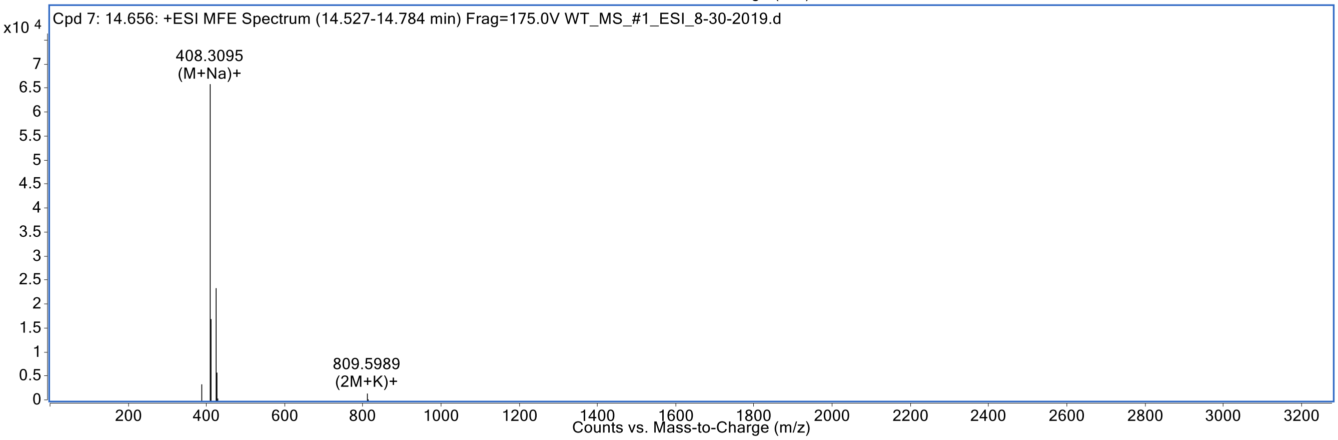
Cpd 7: 14.656: +ESI MFE Spectrum (14.527-14.784 min) Frag=175.0V WT_MS_#1_ESI_8-30-2019.d
x10 4
408.3095
(M+Na)+
809.5989
(2M+K)+
Counts vs. Mass-to-Charge (m/z)

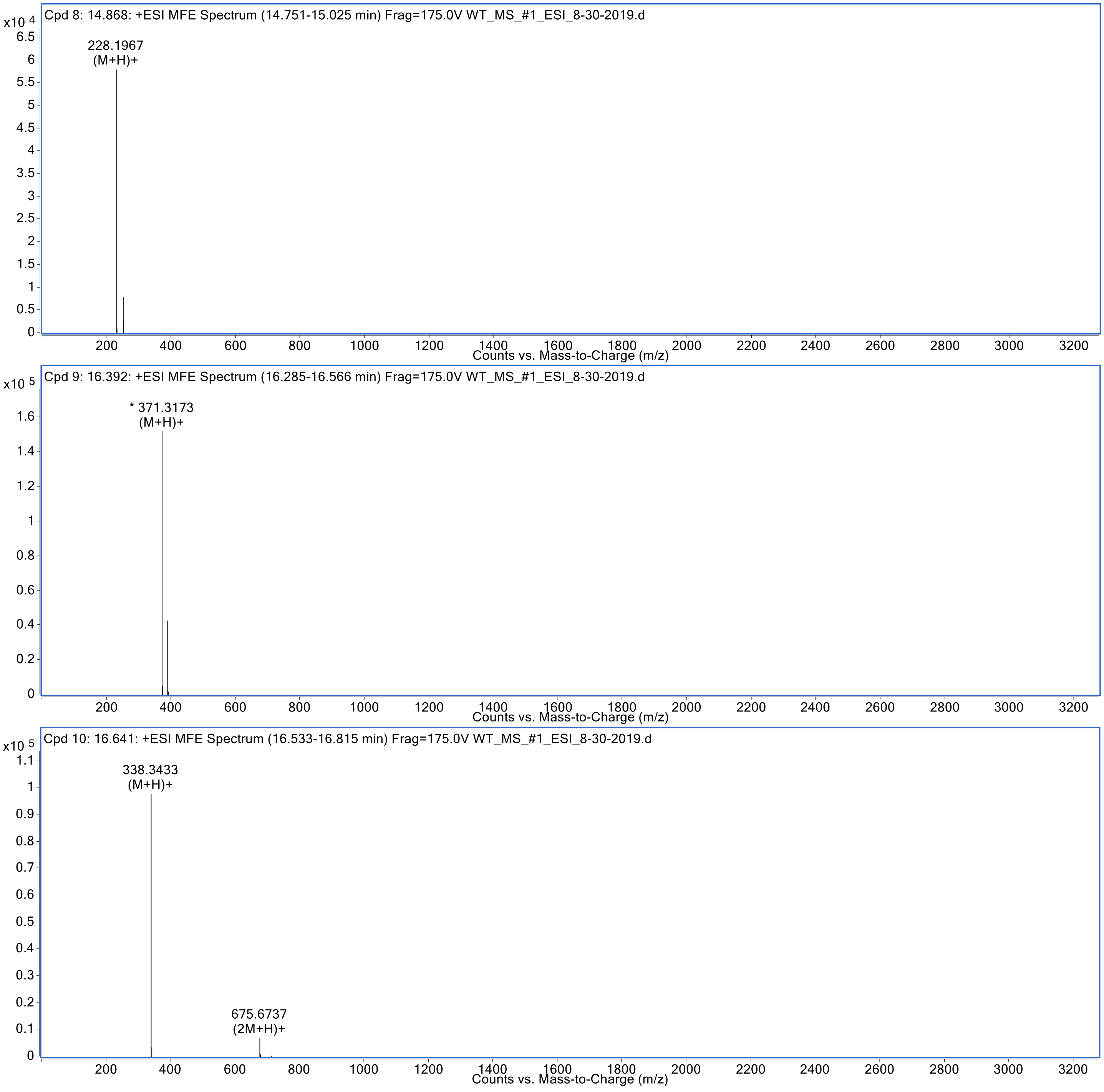


**Figure S4.** LC-MS spectra of 50 μM Aβ(1-40) incubated with 5 μM wild-type IDE (WT_MS_#1).

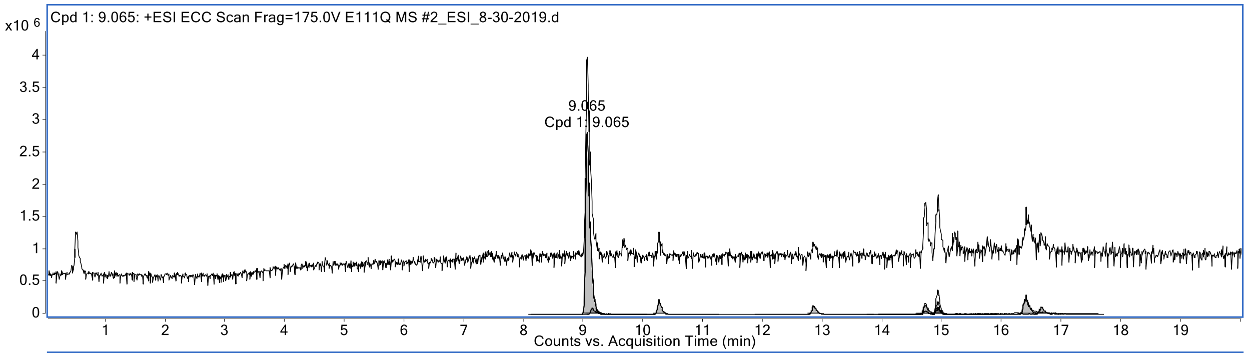
Cpd 1: 9.065: +ESI ECC Scan Frag=175.0V E111Q MS #2_ESI_8-30-2019.d
x10 6
9.065
Cpd 1: 9.065
Counts vs. Acquisition Time (min)

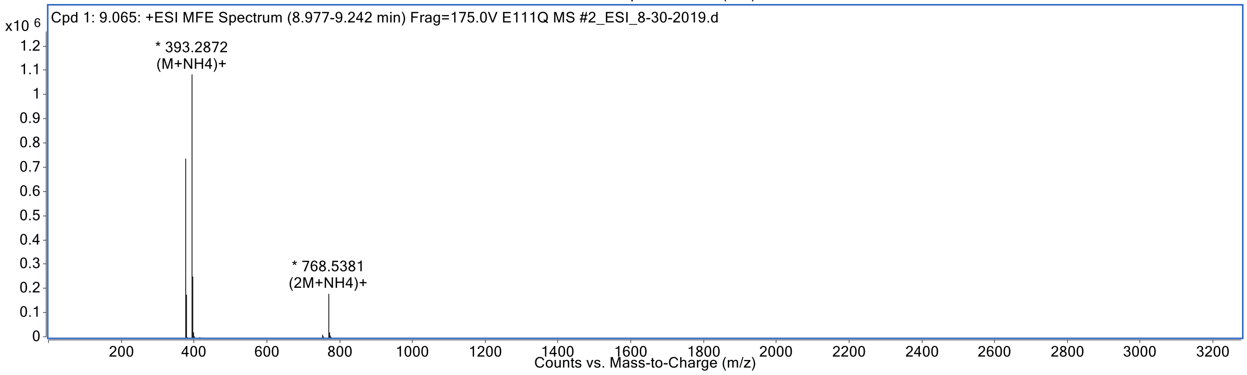
Cpd 1: 9.065: +ESI MFE Spectrum (8.977-9.242 min) Frag=175.0V E111Q MS #2_ESI_8-30-2019.d
x10 6
* 393.2872
(M+NH4)+
* 768.5381
(2M+NH4)+
Counts vs. Mass-to-Charge (m/z)

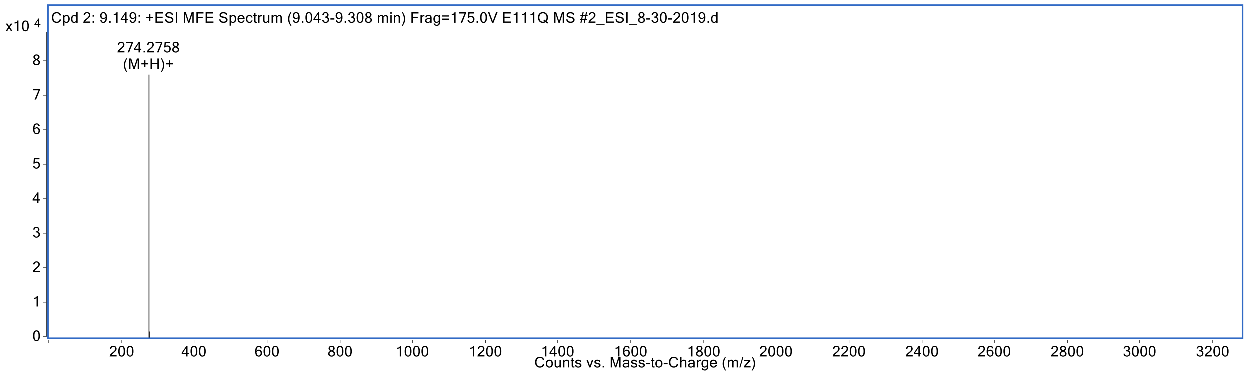
Cpd 2: 9.149: +ESI MFE Spectrum (9.043-9.308 min) Frag=175.0V E111Q MS #2_ESI_8-30-2019.d
x10 4
274.2758
(M+H)+
Counts vs. Mass-to-Charge (m/z)

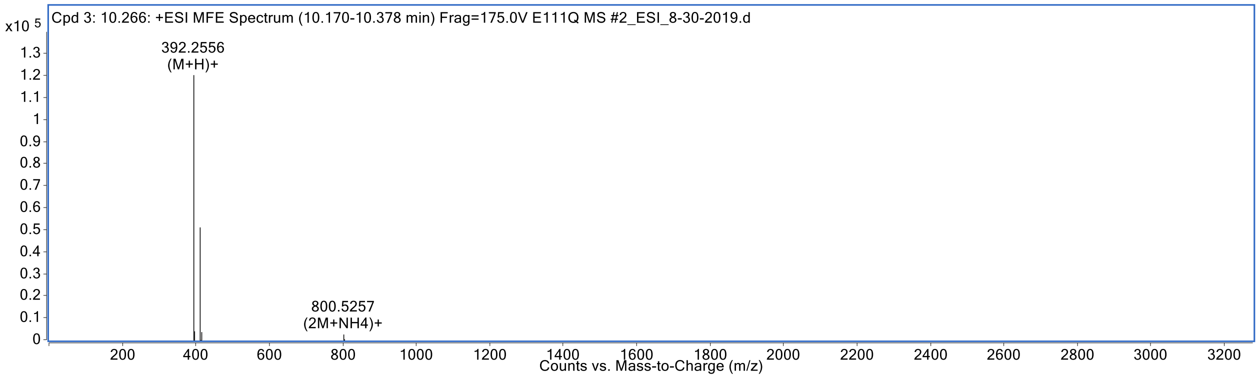
Cpd 3: 10.266: +ESI MFE Spectrum (10.170-10.378 min) Frag=175.0V E111Q MS #2_ESI_8-30-2019.d
x10 5
392.2556
(M+H)+
800.5257
(2M+NH4)+
Counts vs. Mass-to-Charge (m/z)

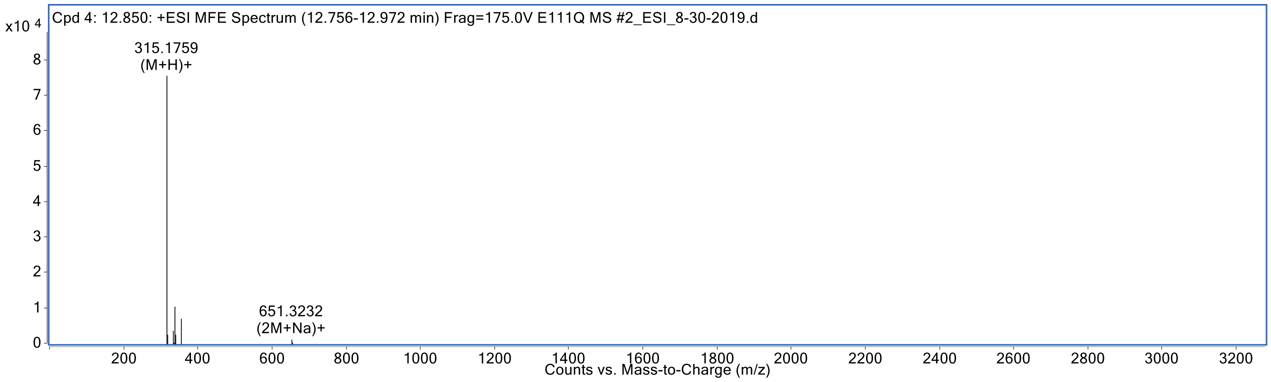
Cpd 4: 12.850: +ESI MFE Spectrum (12.756-12.972 min) Frag=175.0V E111Q MS #2_ESI_8-30-2019.d
x10 4
315.1759
(M+H)+
651.3232
(2M+Na)+
Counts vs. Mass-to-Charge (m/z)

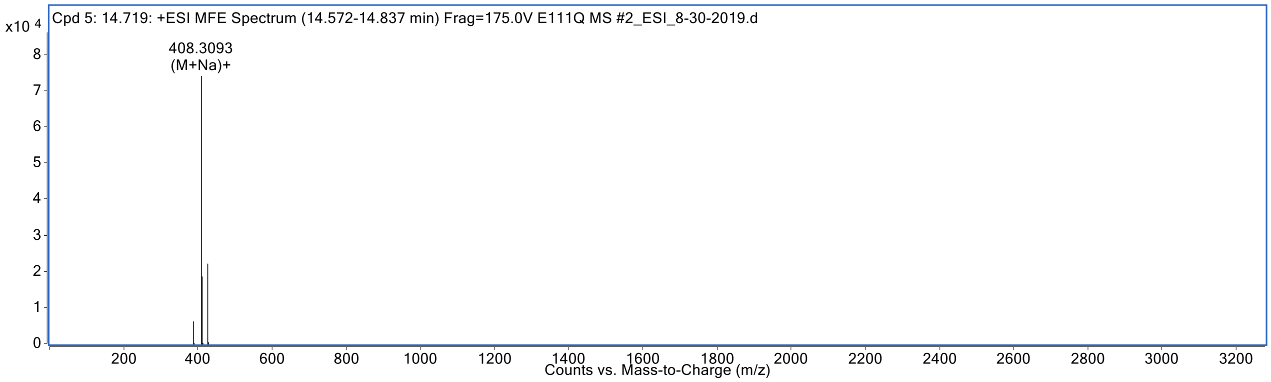
Cpd 5: 14.719: +ESI MFE Spectrum (14.572-14.837 min) Frag=175.0V E111Q MS #2_ESI_8-30-2019.d
x10 4
408.3093
(M+Na)+
Counts vs. Mass-to-Charge (m/z)

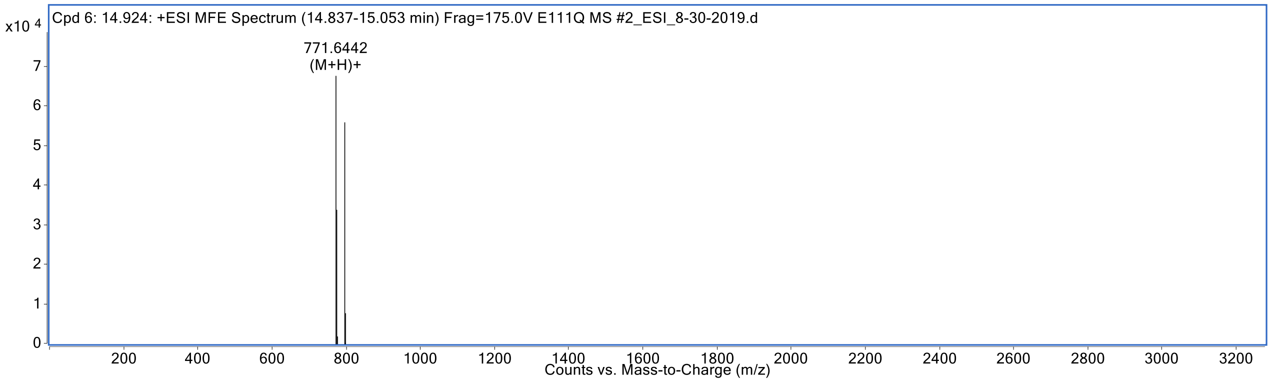
Cpd 6: 14.924: +ESI MFE Spectrum (14.837-15.053 min) Frag=175.0V E111Q MS #2_ESI_8-30-2019.d
x10 4
771.6442
(M+H)+
Counts vs. Mass-to-Charge (m/z)

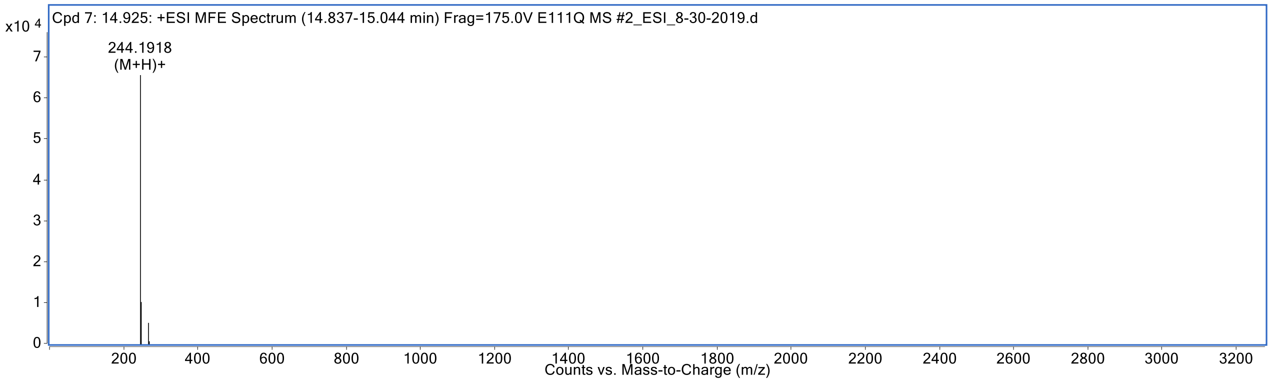
Cpd 7: 14.925: +ESI MFE Spectrum (14.837-15.044 min) Frag=175.0V E111Q MS #2_ESI_8-30-2019.d
x10 4
244.1918
(M+H)+
Counts vs. Mass-to-Charge (m/z)

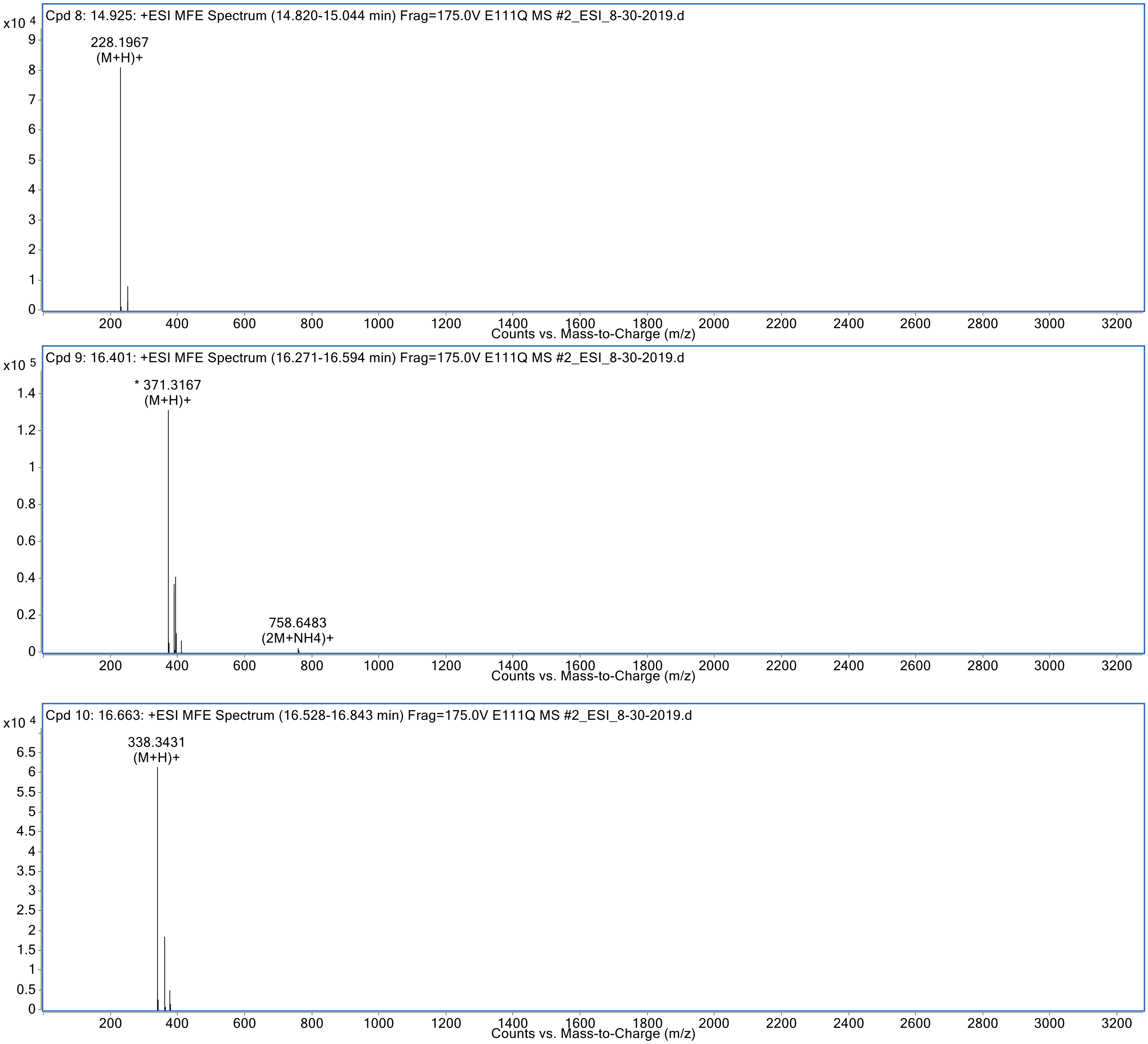


**Figure S5.** LC-MS spectra of 50 μM Aβ(1-40) incubated with 5 μM cysteine-free E111Q-IDE mutant (E111Q_MS_#2).

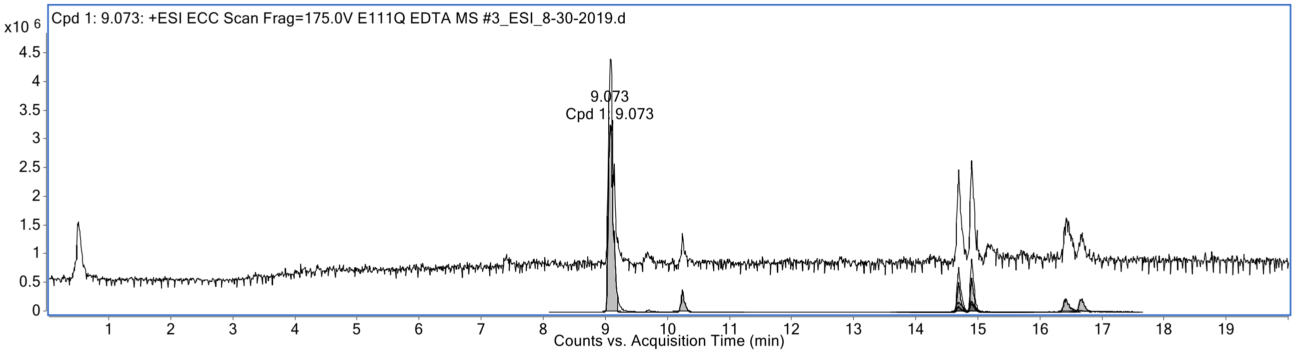
Cpd 1: 9.073: +ESI ECC Scan Frag=175.0V E111Q EDTA MS #3_ESI_8-30-2019.d
x10 6
9.073
Cpd 1: 9.073
Counts vs. Acquisition Time (min)

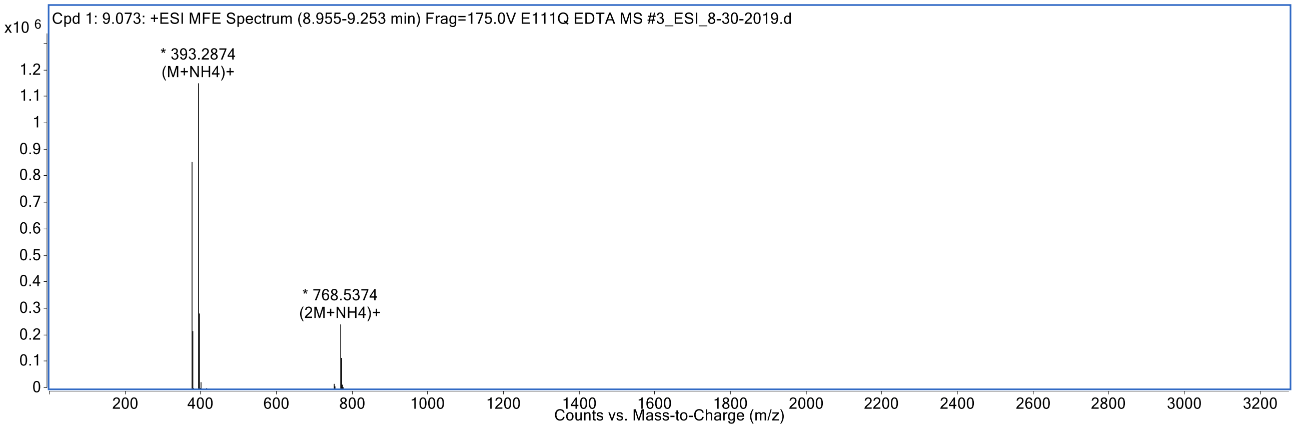
Cpd 1: 9.073: +ESI MFE Spectrum (8.955-9.253 min) Frag=175.0V E111Q EDTA MS #3_ESI_8-30-2019.d
x10 6
* 393.2874
(M+NH4)+
* 768.5374
(2M+NH4)+
Counts vs. Mass-to-Charge (m/z)

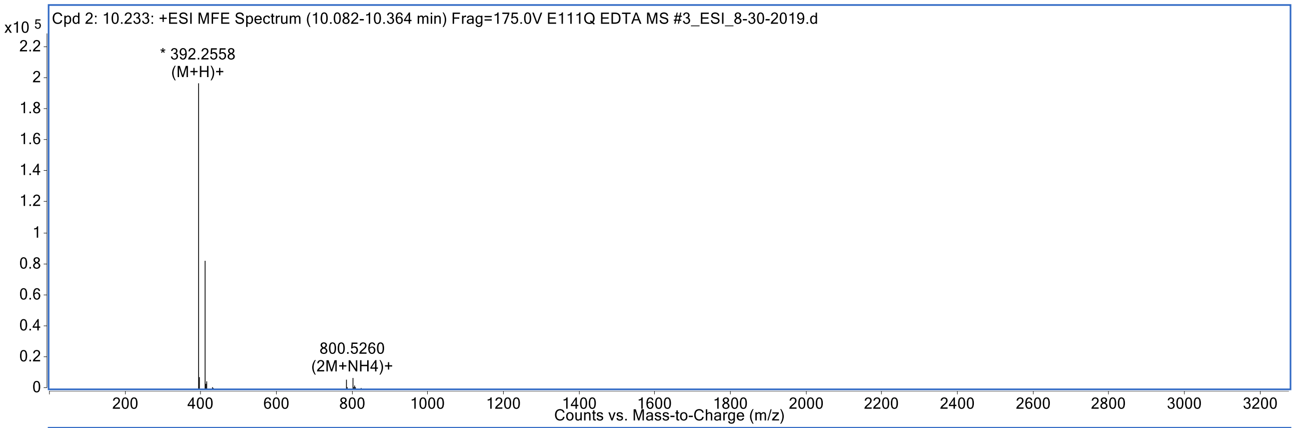
Cpd 2: 10.233: +ESI MFE Spectrum (10.082-10.364 min) Frag=175.0V E111Q EDTA MS #3_ESI_8-30-2019.d
x10 5
* 392.2558
(M+H)+
800.5260
(2M+NH4)+
Counts vs. Mass-to-Charge (m/z)

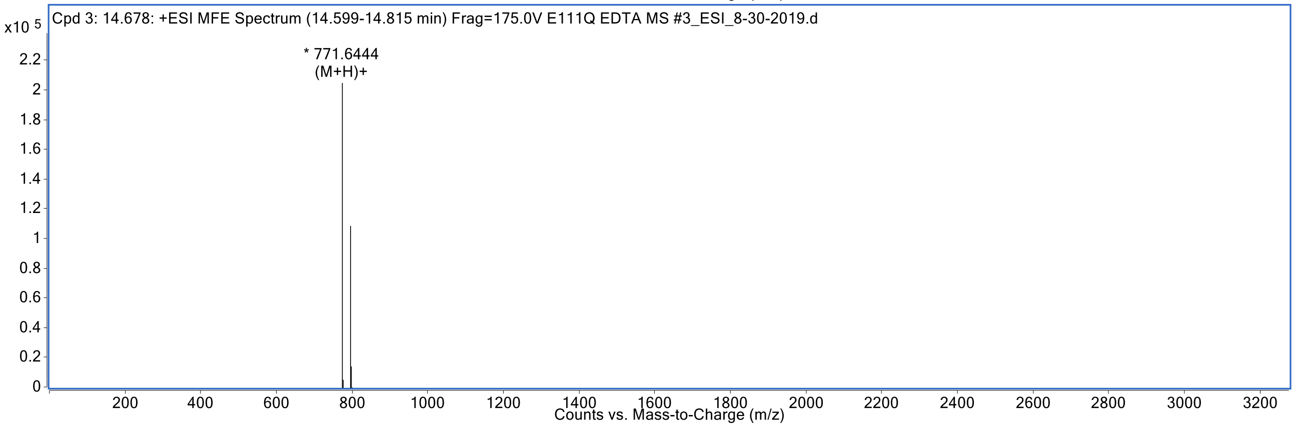
Cpd 3: 14.678: +ESI MFE Spectrum (14.599-14.815 min) Frag=175.0V E111Q EDTA MS #3_ESI_8-30-2019.d
x10 5
* 771.6444
(M+H)+
Counts vs. Mass-to-Charge (m/z)

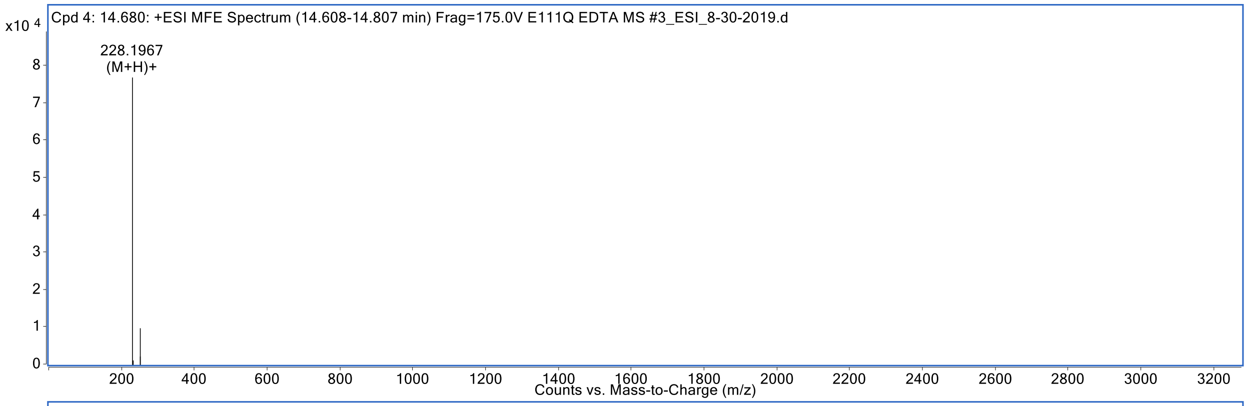

Cpd 4: 14.680: +ESI MFE Spectrum (14.608-14.807 min) Frag=175.0V E111Q EDTA MS #3_ESI_8-30-2019.d
x10 4
228.1967
(M+H)+
Counts vs. Mass-to-Charge (m/z)


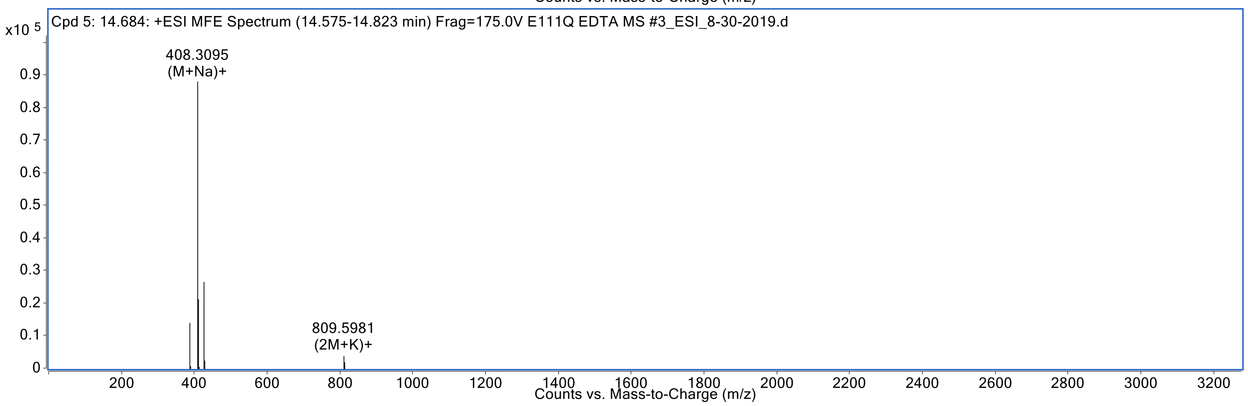

Cpd 5: 14.684: +ESI MFE Spectrum (14.575-14.823 min) Frag=175.0V E111Q EDTA MS #3_ESI_8-30-2019.d
x10 5
408.3095
(M+Na)+
809.5981
(2M+K)+
Counts vs. Mass-to-Charge (m/z)


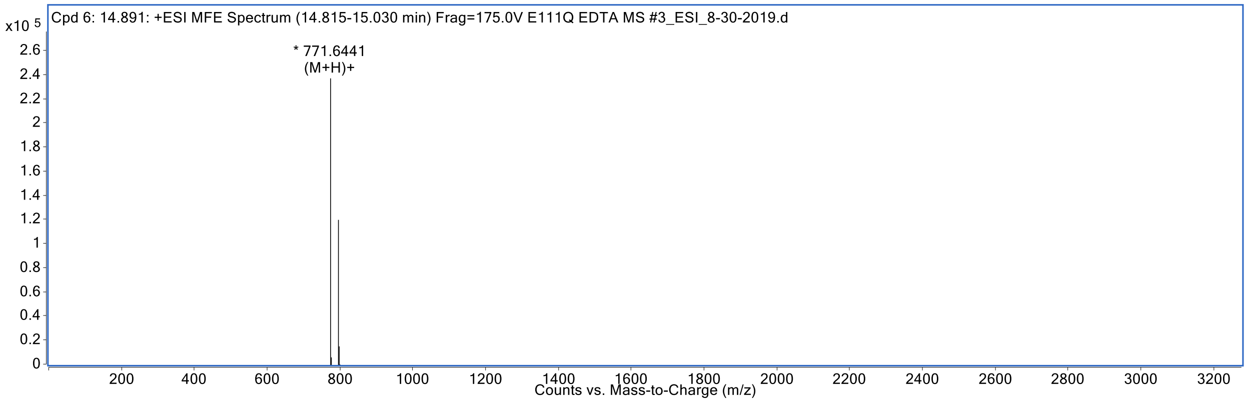

Cpd 6: 14.891: +ESI MFE Spectrum (14.815-15.030 min) Frag=175.0V E111Q EDTA MS #3_ESI_8-30-2019.d
x10 5
* 771.6441
(M+H)+
Counts vs. Mass-to-Charge (m/z)


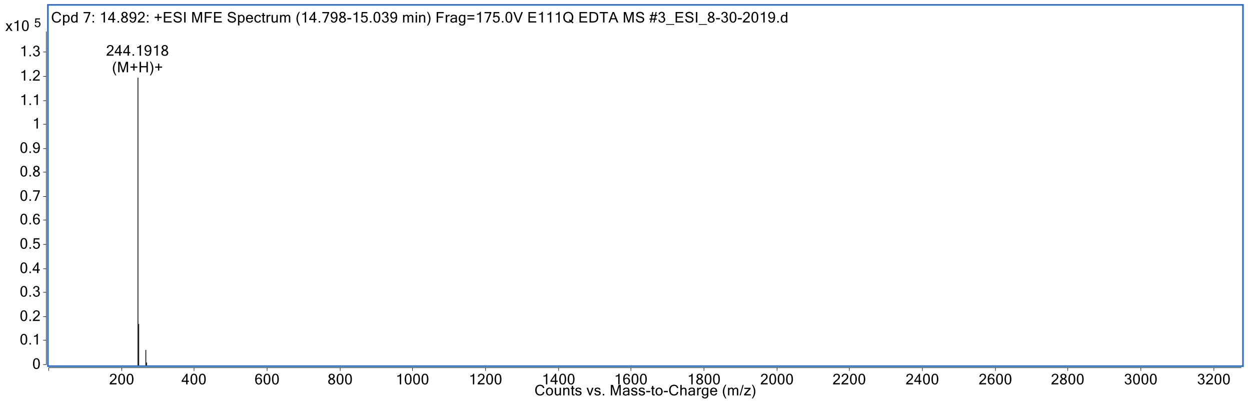

Cpd 7: 14.892: +ESI MFE Spectrum (14.798-15.039 min) Frag=175.0V E111Q EDTA MS #3_ESI_8-30-2019.d
x10 5
244.1918
(M+H)+
Counts vs. Mass-to-Charge (m/z)

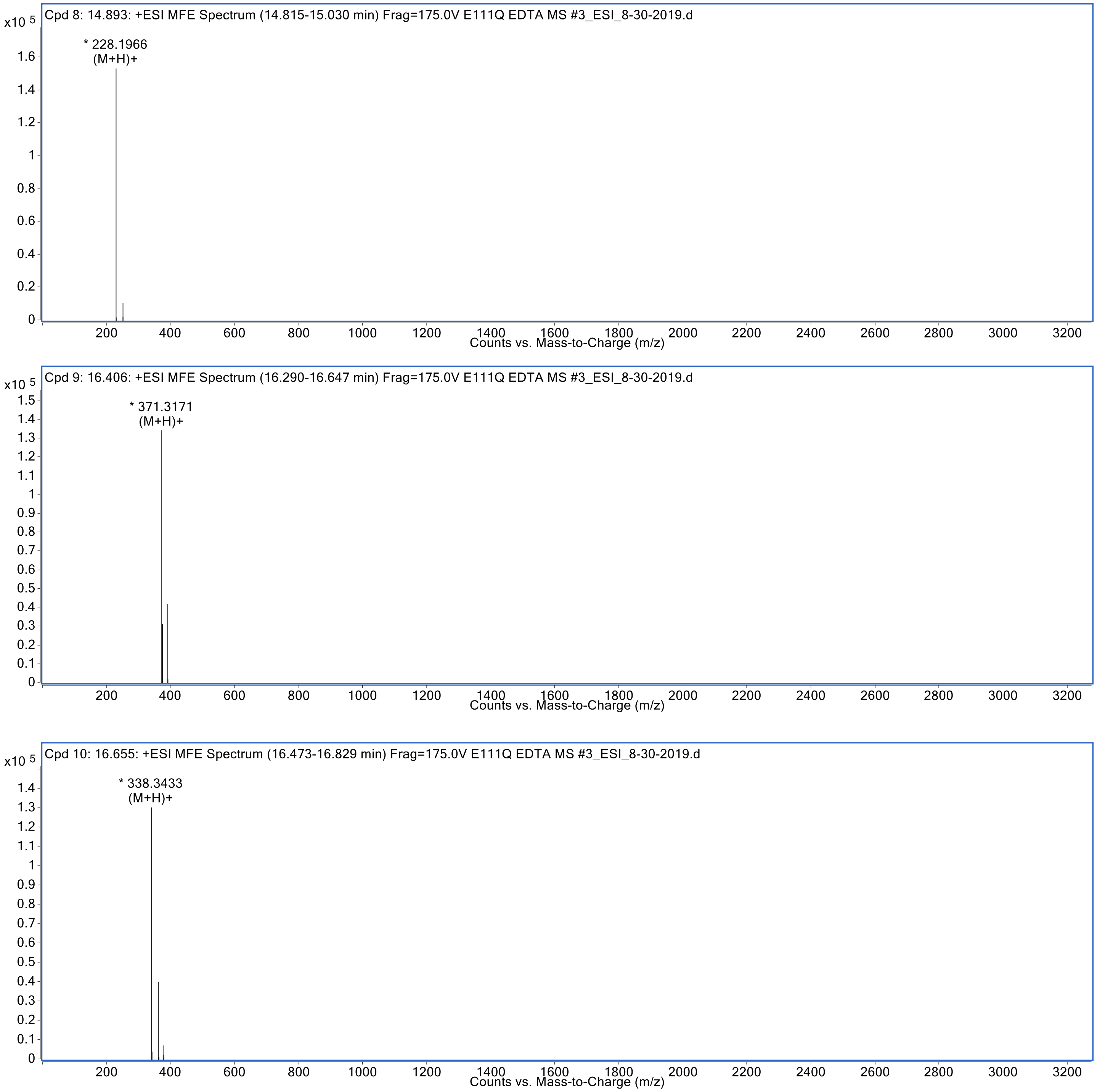


**Figure S6.** LC-MS spectra of 50 μM Aβ(1-40) co-incubated with 5 μM cysteine-free E111Q-IDE + 50 μM EDTA (E111Q_EDTA_MS_#3).

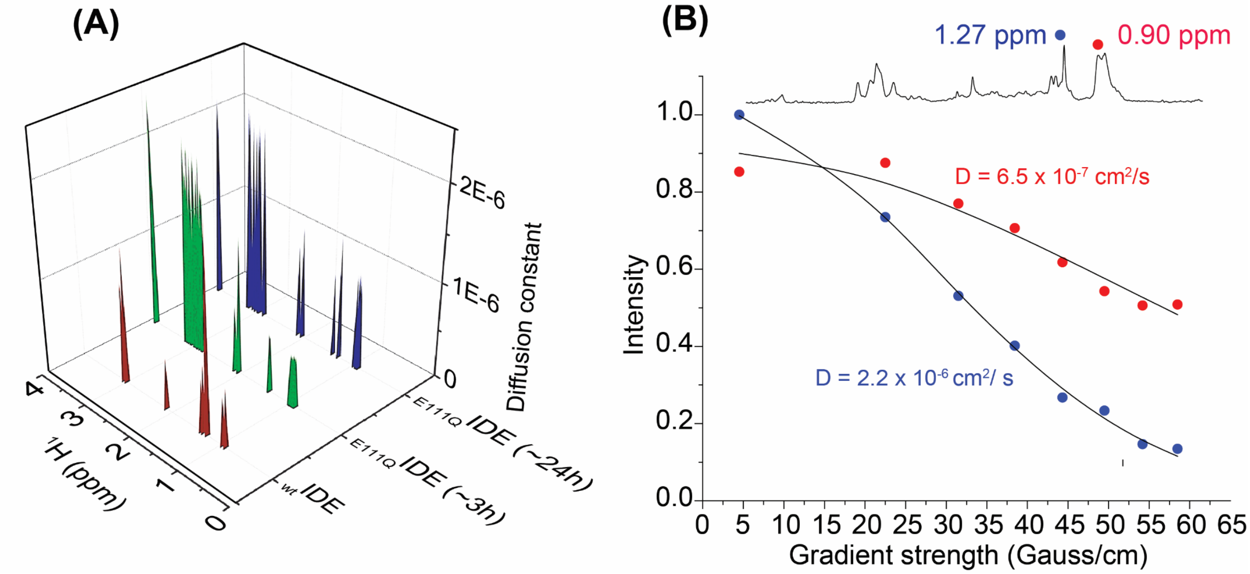


**Figure S7.** (A) 3D plot showing the diffusion constant (cm$^2$/s) of 25 µM Aβ(1-40) fragments cleaved by 5 µM cysteine-free-E111Q-IDE at the indicated time-intervals. (B) Graph showing the diffusion constant, D, values measured from the two representative $^1$H NMR peaks at 0.90 and 1.27 ppm of 25 µM Aβ(1-40) incubated with 2.5 µM of cysteine-free-E111Q-IDE. The DOSY NMR spectra were recorded on a 500 MHz Bruker NMR spectrometer at 25 °C in 10 mM NaPi, pH 7.4 containing 10% $D_2O$, and the corresponding 2D spectra are shown in the main text in Figure 2(E and F).

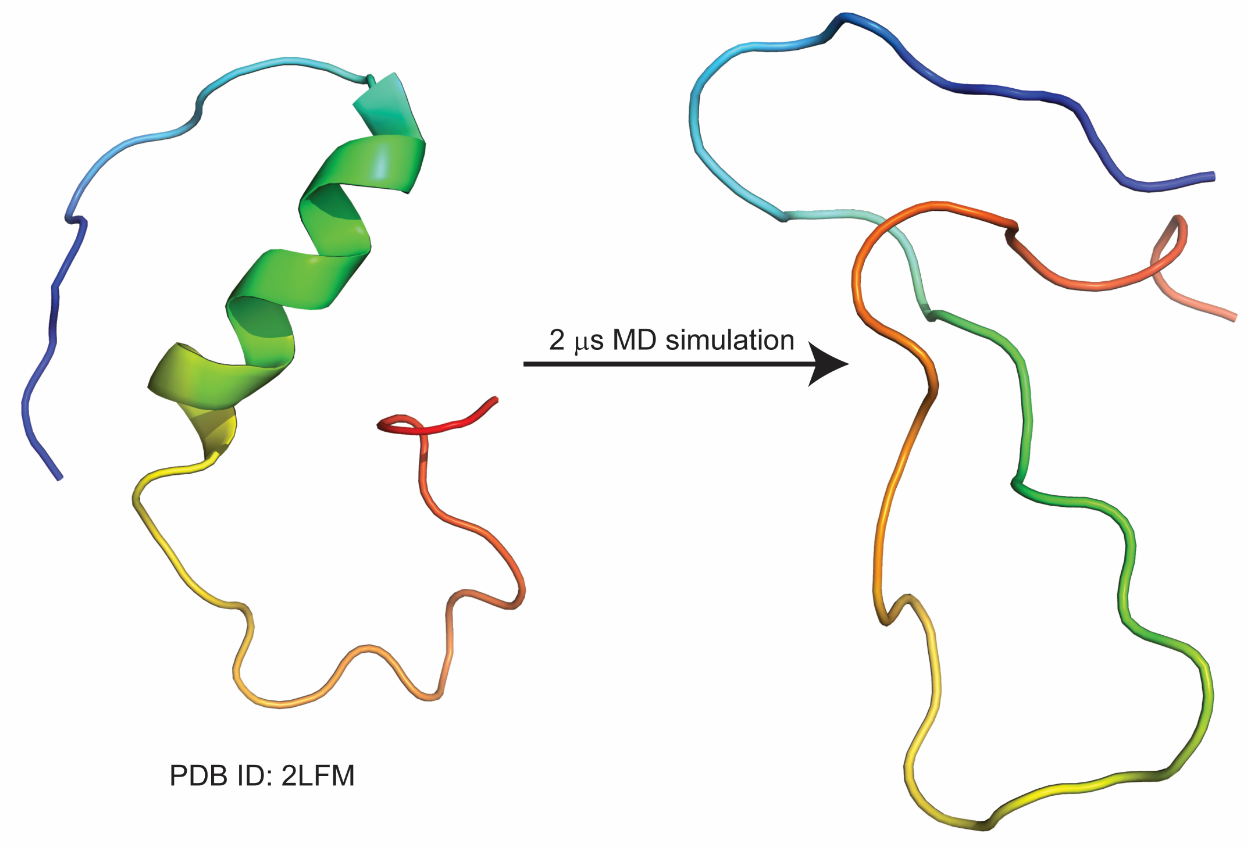


**Figure S8.** A cartoon representation of the misfolded Aβ(1-40) structure (right) obtained after 2 μs MD simulation of the NMR structure (PDB ID: 2LFM) in an aqueous environment. The Aβ(1-40) structure shown on the right was used in this study to generate the initial structures of the 20 different Aβ fragments for MD calculation as shown in the main text Figure 3A.

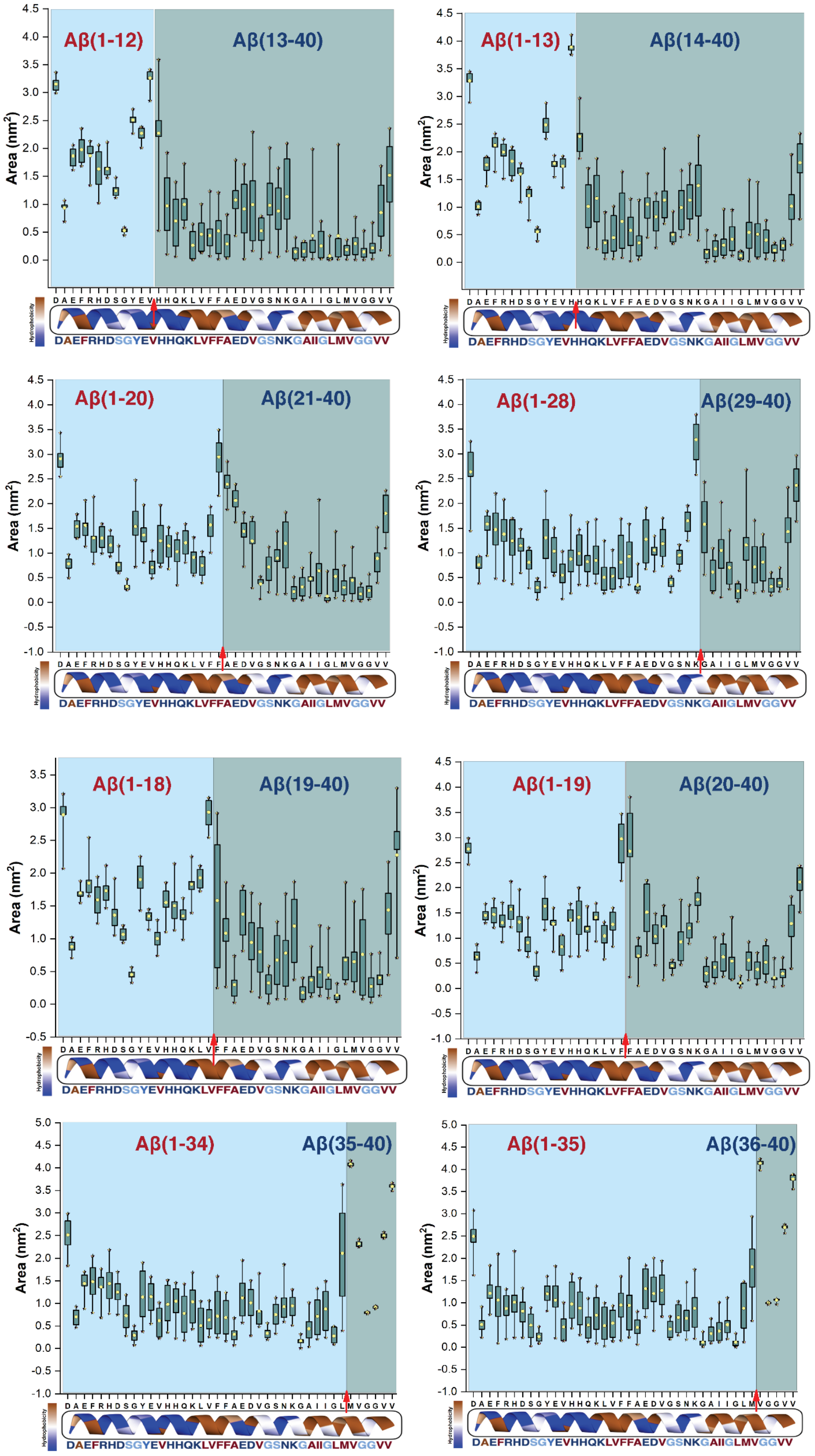


**Figure S9.** Residue-specific average area of the full-length and different fragments of Aβ as indicated. The full length Aβ(1-40) residues presented an average residue-specific area of ≤1.0 $nm^2$, indicating self-assembling during the 500 ns MD simulation.

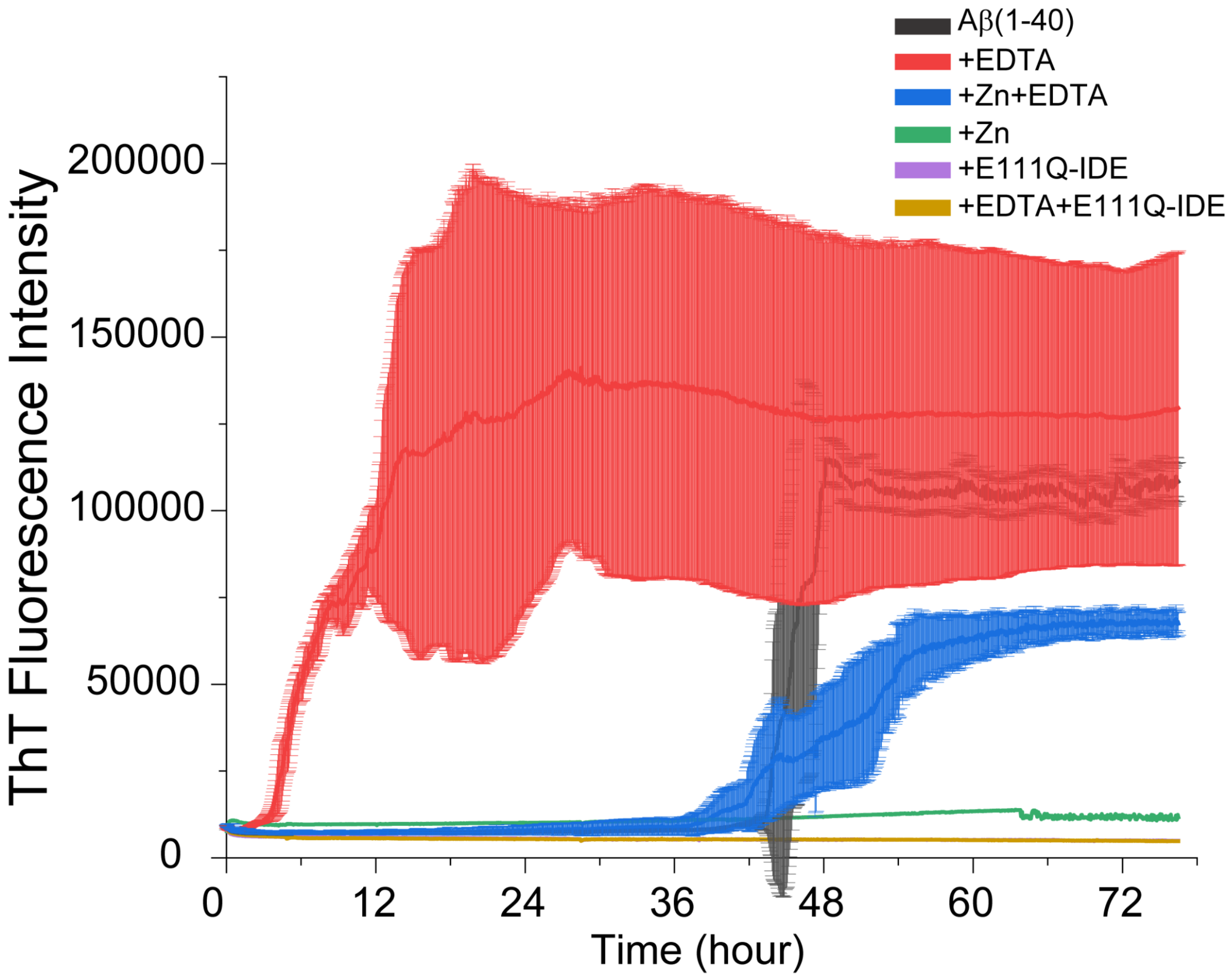


**Figure S10.** Aggregation kinetics of 5 μM Aβ(1-40) using a ThT fluorescence assay in the absence and presence of 5 μM zinc, EDTA and 0.5 μM cysteine-free-E111Q-IDE mutant at the indicated colors (a figure with no error bar is shown in the main text Figure 5A). The fluorescence measurements were carried out in triplicate at 25 °C under no-shaking conditions. The average ThT curve is shown in solid and the standard deviation is shown in shaded color bars.

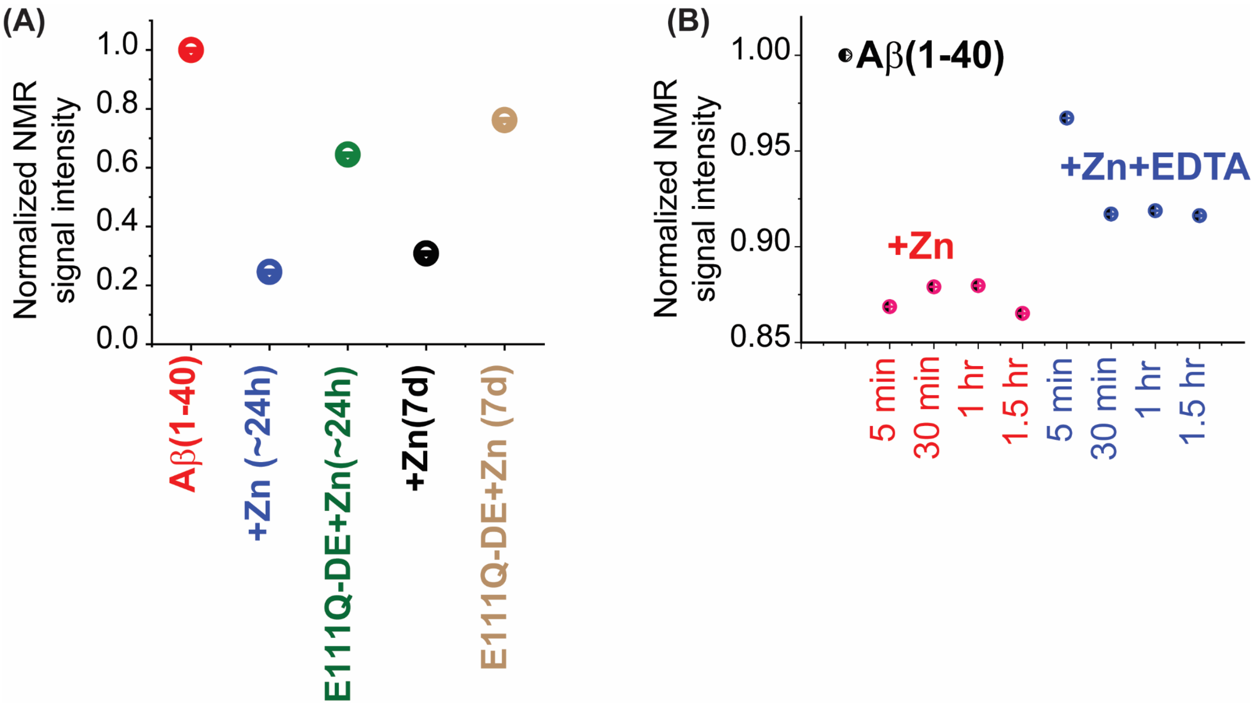


**Figure S11**. (A) Decay of $^{1}H$ NMR signal intensity of 25 μM Aβ(1-40) co-incubated with 25 μM zinc in the absence and presence of 2.5 μM E111Q-IDE at the indicated times. The corresponding 1D NMR spectra are shown in Figure 5G. The NMR signal intensities in the sample containing E111Q-IDE have contributions from both Aβ(1-40) and IDE. (B) $^{1}H$ NMR signal intensity of 25μM Aβ(1-40) showing a reversible change upon zinc (25 μM) binding and zinc removal using 25 μM EDTA. The change in signal intensities are normalized with respect to Aβ(1-40) in the absence of any additives. The corresponding NMR spectra are shown in Figure 5B in the main text. All NMR samples were prepared using 10 mM NaPi buffer, pH=7.4 containing 10% $D_2O$, and spectra were recorded on a 500 MHz Bruker NMR spectrometer at 25 °C.

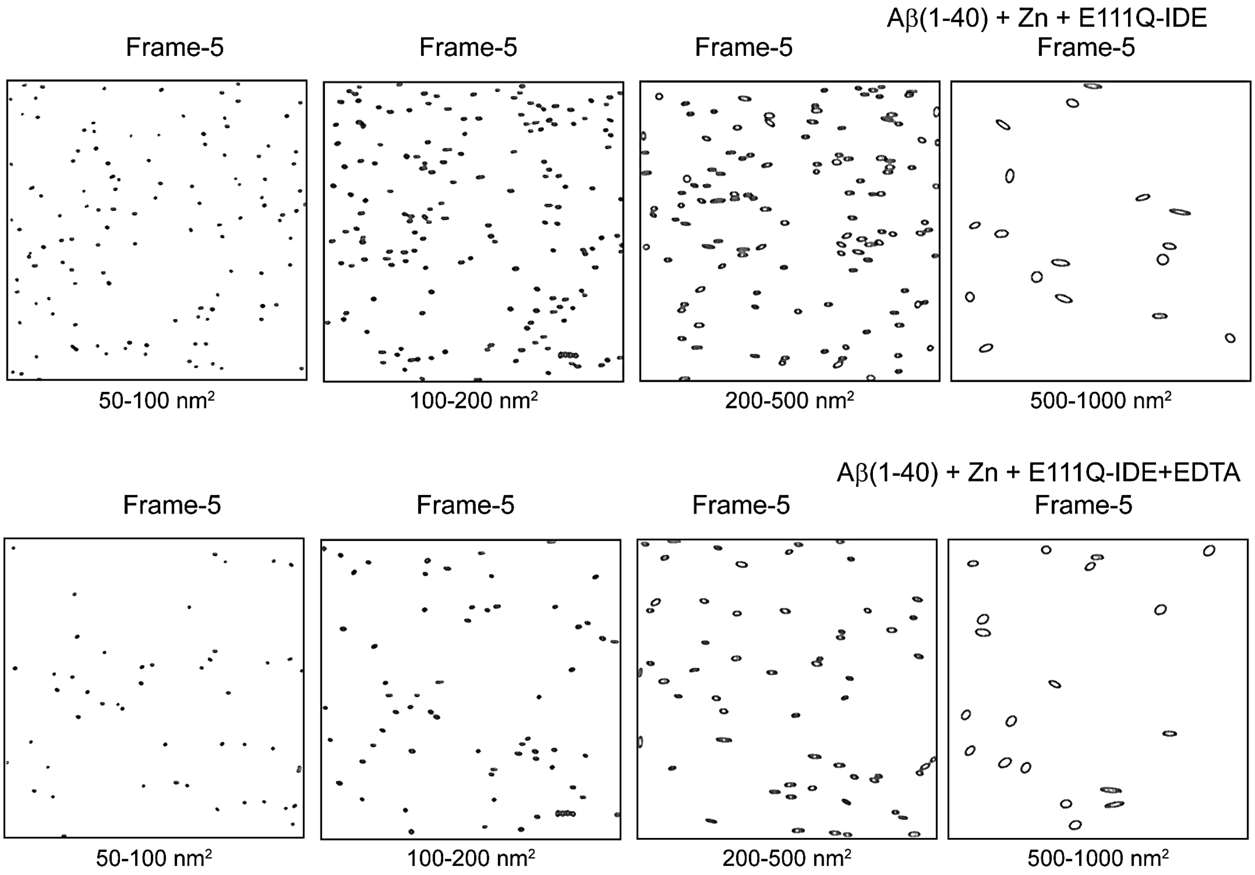


**Figure S12.** Plots for the size distribution analysis of a sample mixture containing 5 μM Aβ(1-40), 5 μM zinc, and 0.5 μM cysteine-free-E111Q-IDE mutant in the absence and presence of 5 μM EDTA as indicated. The particle size was derived from the HS-AFM images frame-5 shown in Figure 6A and 6C using ImageJ. The size-filter used to retrieve the particles are indicated under each plot.

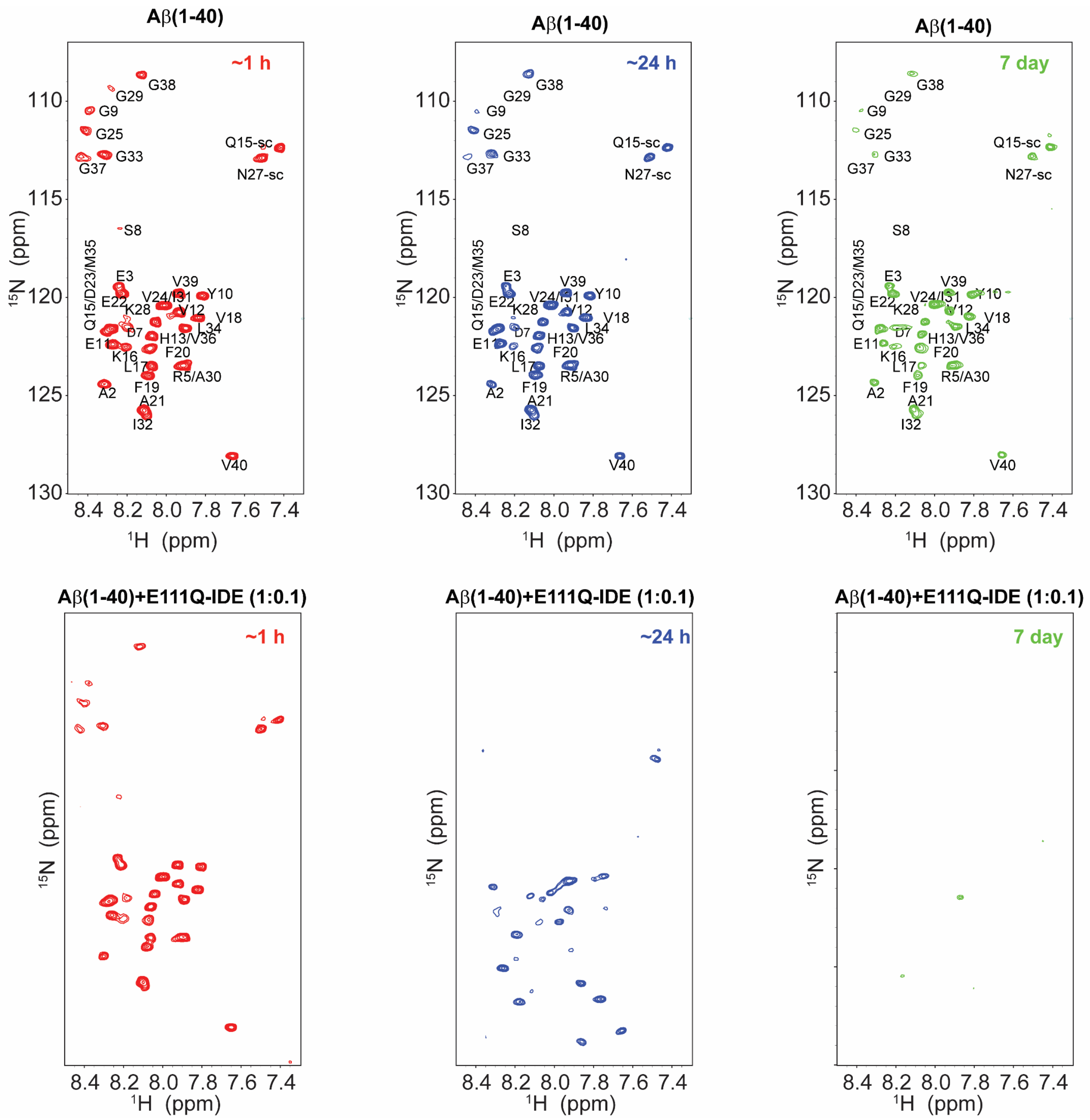


**Figure S13.** Extended figure showing the non-overlapping spectra of Aβ(1-40) and Aβ(1-40) mixed with the cf-E111Q-IDE mutant at 1:0.1 (peptide:enzyme) molar ratio. The corresponding overlapped spectra are shown in the main text Figure 1(I and J).

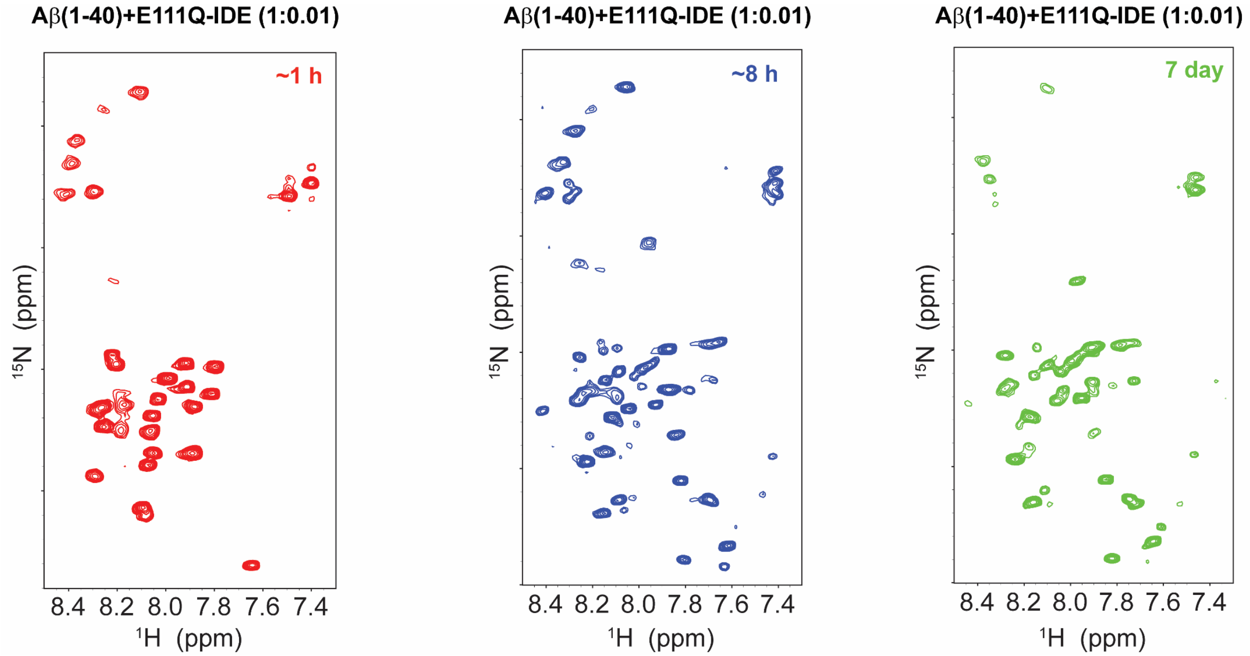


**Figure S14.** Extended figure showing the non-overlapping spectra of Aβ(1-40) mixed with the cf-E111Q-IDE mutant at 1:0.01 (peptide:enzyme) molar ratio. The corresponding overlapped spectra are shown in the main text Figure 1(K and L).

### 4. Supporting videos

4.1 **SV1:** Real-time monitoring of the zinc-Aβ40 complex and the growth in molecular size through association in the presence of E111Q-IDE using HS-AFM.

4.2 **SV2:** Real-time monitoring of the zinc-Aβ40 complex dissociation in the presence of E111Q-IDE titrated with EDTA using HS-AFM.
